# Diffuse ions coordinate dynamics in a ribonucleoprotein assembly

**DOI:** 10.1101/2021.06.25.448160

**Authors:** Ailun Wang, Mariana Levi, Udayan Mohanty, Paul C. Whitford

## Abstract

Proper ionic concentrations are required for the functional dynamics of RNA and ribonucleoprotein (RNP) assemblies. While experimental and computational techniques have provided many insights into the properties of chelated ions, less is known about the energetic contributions of diffuse ions to large-scale conformational rearrangements. To address this, we present a model that is designed to quantify the influence of diffuse monovalent and divalent ions on the dynamics of biomolecular assemblies. This model employs all-atom (non-H) resolution and explicit ions, where effective potentials account for hydration effects. We first show that the model accurately predicts the number of excess Mg^2+^ ions for prototypical RNA systems, at a level comparable to modern coarse-grained models. We then apply the model to a complete ribosome and show how the balance between diffuse Mg^2+^ and K^+^ ions can control the dynamics of tRNA molecules during translation. The model predicts differential effects of diffuse ions on the free-energy barrier associated with tRNA entry and the energy of tRNA binding to the ribosome. Together, this analysis reveals the direct impact of diffuse ions on the dynamics of an RNP assembly.

**TOC Graphic:** 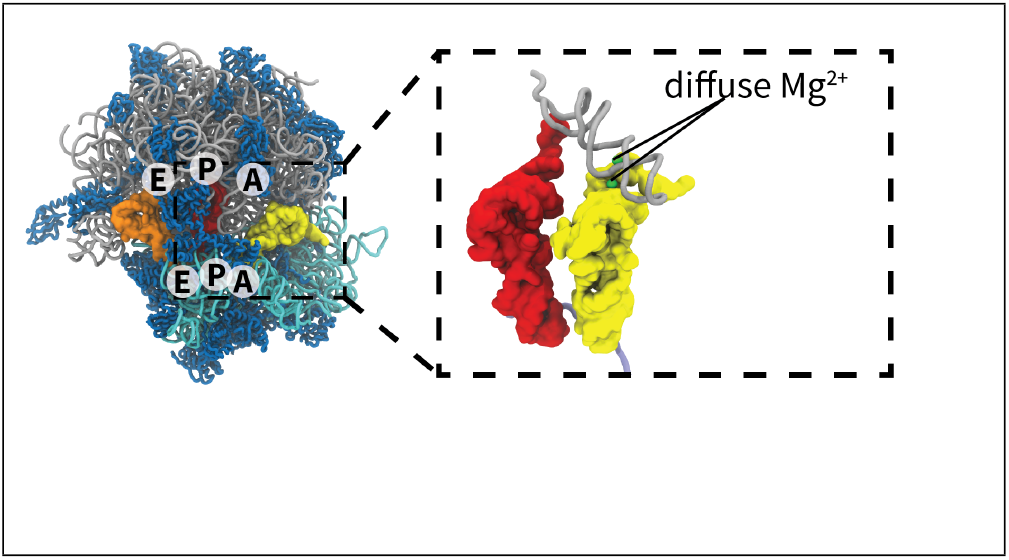

## Introduction

Many large-scale biomolecular processes in the cell depend on the presence of monovalent and multivalent ions. The contribution of cations to structure and dynamics has been experimentally documented for a variety of systems, including RNA,^1–6^ DNA^7,8^ and ribonucleoprotein (RNP) assemblies.^9,10^ A particularly well-characterized RNP assembly is the ribosome, for which specific counterion concentrations are needed for assembly^11,12^ and conformational transitions between functional states.^13,14^ In terms of biological function, *in vitro* studies have revealed how ions can even regulate the accuracy of protein synthesis by the ribosome.^15,16^ While the broad influence of counterions is acknowledged, identifying the physical-chemical relationship between ions and the mechanical properties of large-scale RNP assemblies has remained elusive.

The solvent environment around RNA is generally described as containing chelated and diffuse ions.^3,4^ Chelated (i.e. inner-shell) ions are partially dehydrated, which allows them to form strong direct contacts with RNA.^3,4,17^ As a result, chelated ions can remain bound to RNA for ms-s timescales, ^18–20^ which exceeds the duration of many biomolecular processes. In contrast, diffuse (outer-shell) ions maintain a coordinated hydration shell (e.g. 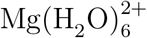) and associate less strongly with RNA. Even though water molecules can exchange rapidly (microseconds), ^21–23^ each ion remains fully hydrated. Accordingly, the behavior of a diffuse ion is primarily determined by longer-range electrostatic interactions. Despite the transient and weak influence of individual diffuse ions, their collective effect can be significant. For example, tRNA structural stability increases with the concentration of monovalent cations, which can be attributed to screening effects and ion-mediated interactions.^24,25^ The diffuse ionic environment can also control tertiary structure formation in RNA,^3^ including ribozymes^26^ and the ribosome.^27^ A complicating factor when studying diffuse ions is that monovalent and divalent ions competitively associate with RNA. Due to this, a balance of entropic and enthalpic factors can lead to non-trivial relationships between ionic concentrations and biomolecular stability/dynamics.

The essential roles of ions in biology has motivated their study over a wide range of temporal and spatial scales. There are now many computational approaches available that include implicit-solvent and all-atom explicit-solvent representations, as well as coarse-grained models. Within the class of implicit-solvent models, there have been applications of non-linear Poisson Boltzmann theory,^28–30^ counterions condensation models,^31–36^ reference interaction site model^37–39^ and Debye-Hückel (DH) treatments,^40–42^ among other strategies.^43–45^ While these methods can provide accurate representations of the local electric fields, it is common for these approaches to neglect ion-ion correlations, which can make it difficult to distinguish between the effects of monovalent and divalent species.^46,47^ In the study of biomolecular assemblies, coarse-grained models frequently employ a DH treatment for monovalent ions. ^48^ This is appropriate for describing interactions between opposing charges (e.g. protein-DNA association^49,50^), though it can not (by construction) capture ion-induced attraction between polyanionic molecules.^51^ To address this shortcoming, coarse-grained models that include explicit-ion representations have been developed to analyze ion-mediated attraction in DNA.^52,53^ Explicit-ion models with coarse-grained RNA representations have also been successful in predicting biomolecular stability,^42,54^ as well as the mechanisms^26,27^ and energetics^39^ of large RNA folding. At a higher level of spatial resolution, ion parameters are available for use with all-atom explicit-solvent simulations.^55–57^ In various applications, these have provided interpretations of ensemble experimental data^58^ and insights into the energetics of small RNA systems.^59–61^ While explicit-solvent techniques may be applied to larger systems, the associated computational requirements have limited the accessible timescales to microseconds. ^62^ As a result, there is not an all-atom model available for which it is tractable to directly connect the properties of diffuse ions with slow (millisecond) conformational processes in large-scale RNP assemblies.

To probe how diffuse ions influence conformational dynamics in biomolecular assemblies, we developed an all-atom (non-Hydrogen atom) model that employs simplified energetics for each biomolecule, along with a transferrable effective potential for explicit monovalent (K^+^, Cl^*−*^) and divalent (Mg^2+^) ions. In this model, which we call SMOG-ion, an all-atom structure-based (SMOG) model^63,64^ is used to define intramolecular interactions, while ionic interactions are assigned non-specific effective potentials and Coulomb electrostatics. The ion parameters were first refined based on comparison with explicit-solvent simulations of small model systems (rRNA helix and protein S6). A subset of the ion-RNA parameters were then further refined through comparison with an experimental measure of the excess ionic atmosphere for a prototypical rRNA fragment. While the parameters were defined based on experiments performed for a single system and concentration, we find the model accurately describes the concentration-dependent properties of the diffuse ionic atmosphere for multiple small RNA molecules. Building upon these benchmarks, we applied the model to simulate a bacterial ribosome in the presence of monovalent and divalent ions. We find that the free-energy barrier of a large-scale (∼ 30Å) conformational rearrangement (i.e. tRNA accommodation) is regulated by the diffuse ionic environment. In addition, the affinity of tRNA for the ribosome shows a clear ionic concentration dependence. Together, these calculations implicate a direct relationship between diffuse ions and the structural dynamics of this large biomolecular assembly.

## Methods

### Structure-based “SMOG” model with explicit ions

The SMOG-ion model is an all-atom structure-based “SMOG” model^63,64^ with explicit electrostatics and an explicit representation of diffuse ions (K^+^, Cl^*−*^ and Mg^2+^). Ionic interactions are defined in terms of Coulomb electrostatics and effective potentials. The potential energy may be described in terms of two components:

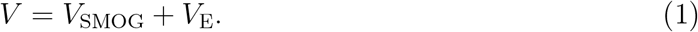

*V*_SMOG_ refers to the all-atom structure-based potential energy and *V*_E_ describes the potential energy of all electrostatic interactions.

In the all-atom structure-based SMOG model (*V*_SMOG_),^63,64^ all non-hydrogen atoms are explicitly represented and an experimentally-identified configuration is defined as the global potential energy minimum. The functional form of the SMOG model is given by:

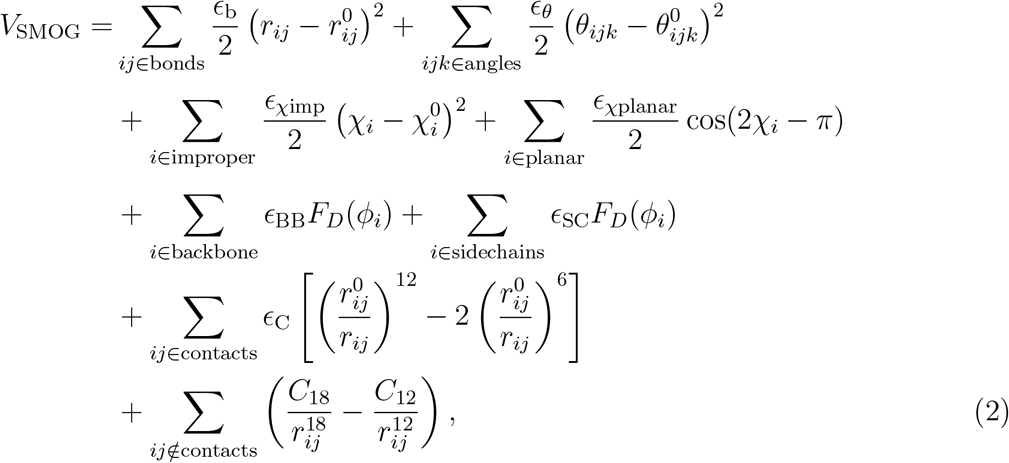

where

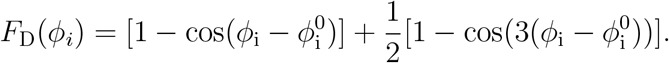

The 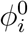 are given the values found in a pre-assigned configuration 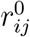 and 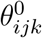 are assigned the corresponding values found in the Amber99sb-ildn force field,^65^ as employed in a previous SMOG-AMBER model. ^66^ Interaction weights are assigned as defined in Ref. 64. Contacts are defined using the Shadow Contact Map algorithm^67^ with a 6Å cutoff and a 1Å shadowing radius.

In contrast with previous SMOG models, we included a 12-18 potential for atom pairs that are not in contact in the experimentally-defined structure. This was introduced in order to define an excluded volume potential that mimics that of the AMBER force field, without including a deep attractive well (Fig. S1). With this approach, the steric representation provided by the model is consistent with the more highly-detailed AMBER model, while introducing a minimal degree of non-specific energetic roughness. The coefficients *C*_18_ and *C*_12_ were defined for each type of interaction based on fits to the corresponding 6-12 potentials in the Amber99sb-ildn force field^65^ (Fig. S1).

The electrostatic representation (*V*_E_) includes direct Coulomb interactions (*V*_coulomb_), effective excluded volume potentials for diffuse ions (*V*_ion*−*excl_) and effective potentials that describe ionic solvation effects (*V*_sol_):

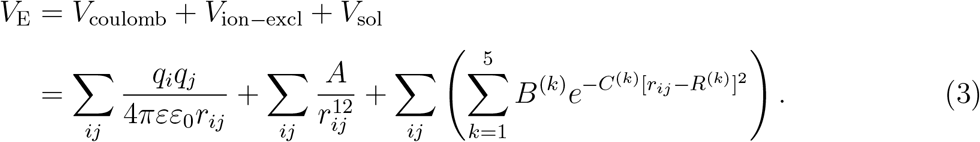

*V*_coulomb_ represents the direct Coulomb interactions between a pair of charges *q*_*i*_ and *q*_*j*_ with interatomic distance *r*_*ij*_. *ε* is the dielectric constant for water (80) and *ε*_0_ is the permittivity of free space. The partial charge of each atom was derived from the Amber99sb-ildn forcefield.^65^ Since the SMOG-ion model only includes non-hydrogen atoms, the partial charge of each hydrogen atom in the AMBER force field was added to the corresponding non-H atom.

The effective excluded volume of each diffuse (hydrated) ion is account for by *V*_ion*−*excl_. Consistent with previous efforts to model explicit ions in a coarse-grained model,^52^ pairwise potentials of the form 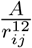 were used to define the excluded volume between ion *i* with atom *j*. The parameter *A* is different for each type of interaction in the model (e.g. Cl-Cl, Cl-K, K-O, etc). To simplify notation, subscripts are not shown. Following the protocol of Savelyev and Papoian,^52^ the values of the parameters *{A}* were obtained through refinement based on comparison with explicit-solvent simulations (See Supporting Information for details).

The term *V*_sol_ describes solvent-mediated ionic interactions, which manifest in the form of ionic shells. The functional form is the same as used previously,^52^ where a sum of Gaussians describe the shells (*B*^(*k*)^ *<* 0) and intervening barriers (*B*^(*k*)^ *>* 0). For each type of interaction considered, up to five Gaussians were included to describe up to three (outer) ionic shells. The location (*R*^(*k*)^) and width (*C*^(*k*)^) of each Gaussian was set based on an initial fit to the corresponding radial distribution function. The amplitude of each Gaussian (*B*^(*k*)^) was refined based on comparison with explicit-solvent simulations and experimental measurements (details in Supporting Information). Consistent with the assignment of excluded volume parameters, unique Gaussian parameters were assigned to each type of modeled interaction. As a note, the parameters in our model are not concentration dependent. Since calculations and experiments have shown the strong concentration dependence of ionic chemical potentials for large changes in ionic strengths (0 *−* 5.5 m),^68^ the parameters in the SMOG-ion model will likely need to be re-refined if one seeks to study higher ionic concentrations that those considered here.

Upon publication, the SMOG-ion model will be freely available through the SMOG 2 Force Field Repository (https://smog-server.org/smog2 Force Field ID: AA_ions_Wang22.v1).

### Calculating preferential interaction coefficients

We calculated the preferential interaction coefficient of Mg^2+^ ions (Γ_2+_) from simulations with the SMOG-ion model and used it as a metric to compare with experimental measurements. Γ_2+_ describes the “excess” number of Mg^2+^ ions that accumulate around an RNA molecule due to electrostatic interactions. ^69–73^ In the current study, Γ_2+_ was calculated by taking the difference between the total number of simulated ions 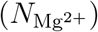 and the expected bulk value. The expected bulk number is the product of the bulk density 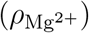 and the volume of the simulated box (*V*_box_)^i^, such that 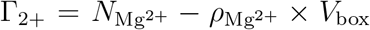. To calculate the bulk density, the RNA fragment was first recentered in the box for each frame of the trajectory. The box was then partitioned into five equal-width (*∼* 140Å) slabs. The average density of Mg^2+^ was then calculated by excluding the central (RNA-containing) slab. For each system and ion concentration, the effective concentration was calculated for 20 independent replicas. The uncertainty in the concentration was defined as the standard deviation of the 20 replicate values. While simulating many replicas helped reduce the uncertainty (standard deviation of [MgCl_2_] *∼* 0.005 mM), due to the large dimensions of the simulated systems, this leads to a residual uncertainty of *∼*1 in *ρ*_Mg2+_ *× V*_box_. This uncertainty is then propagated to Γ_2+_.

When calculating Γ_2+_, it was necessary to account for chelated ions, as well as the diffuse ionic environment. Since the SMOG-ion model was only parameterized to describe diffuse ions, strongly-bound chelated (inner-shell) ions were assigned *a priori*. In the adenine riboswitch, there are five Mg^2+^ binding sites (Fig. S2b) identified in the crystal structure (PDB: 1Y26).^74^ However, the binding site of Mg3 is formed by crystallographic interactions^74^ and no divalent cation binding could be detected in the vicinity of this position through highresolution NMR spectroscopy and titration methods. ^75^ Accordingly, this chelated ion was not included in the current simulations. However, since the other four Mg^2+^ ions are found deep within the grooves of the RNA, they were defined to be harmonically restrained to their chelation pockets. To compare with experimental approaches, which report the number of diffuse and bound ions, the four chelated ions were included in our calculation of Γ_2+_. Chelated ions in the 58-mer are described in the results section.

## Results

To identify the effects of diffuse ions on the dynamics of biomolecular assemblies, we developed an all-atom model with simplified energetics and explicit ions (K^+^, Cl^*−*^, Mg^2+^), which we call SMOG-ion. An all-atom structure-based (SMOG) model^63,64^ with explicit charges defines the biomolecular energetics. Ionic interactions are defined by Coulomb electrostatics and effective potentials (*V*_E_, Eq. 3). The “structure-based” terms explicitly favor a pre-defined biomolecular structure, while the effective potentials ensure that the local distribution of ionic species is consistent with *in vitro* measurements. After describing performance benchmarks and parameterization, we demonstrate how this model may be used to isolate the influence of diffuse ions in a molecular assembly. Specifically, we compare the dynamics of the ribosome with multiple variants of the model, which reveals the direct impact of diffuse ions on the kinetics of a large-scale conformational transition that is central to protein synthesis.

### Simplified model reproduces *in vitro* ionic distribution

To parameterize the SMOG-ion model, we first refined the interaction weights based on comparisons with explicit-solvent simulations of multiple systems (Fig. 1 and Fig. S3) for a single ionic composition ([MgCl_2_] *≈* 10 mM, [KCl] *≈* 100 mM). The explicit-solvent simulations used the Amber99sb-ildn^65^ force field with Mg^2+^ parameters described by Åqvist^55^ and monovalent ion (K^+^ and Cl^*−*^) parameters of Joung and Cheatham.^56^ We refined the ion-ion, ion-RNA and ion-protein interactions in our model using the procedure of Savelyev and Papoian,^52^ where linear parameters in the Hamiltonian are iteratively updated through use of a first-order expansion of the partition function (Eq. S2b). With regards to nomenclature, “s*N* ” denotes the model parameters after refinement <u>s</u>tep *<u>N</u>*. The initial parameters (s0; Fig. 1d, blue curve) were estimated based on inspection of radial distribution functions (RDFs) calculated from explicit-solvent simulations (Fig. S4a). With the s0 parameters, the positions and widths of the peaks in each RDF were consistent with the explicit-solvent model (Fig. 1e, blue curve), though there were significant differences in the heights of each peak. We then sequentially refined the ion-ion, ion-RNA and ion-protein interactions (i.e. s1 parameter set; details in Supporting Information). The s1 model produced RDFs for Mg^2+^, K^+^ and Cl^*−*^ that exhibited excellent agreement with those obtained with the explicit-solvent model (Figs. 1e and S5). This initial refinement ensures the description of outer ionic shells is comparable to predictions by more highly-detailed models.

**Figure 1:**
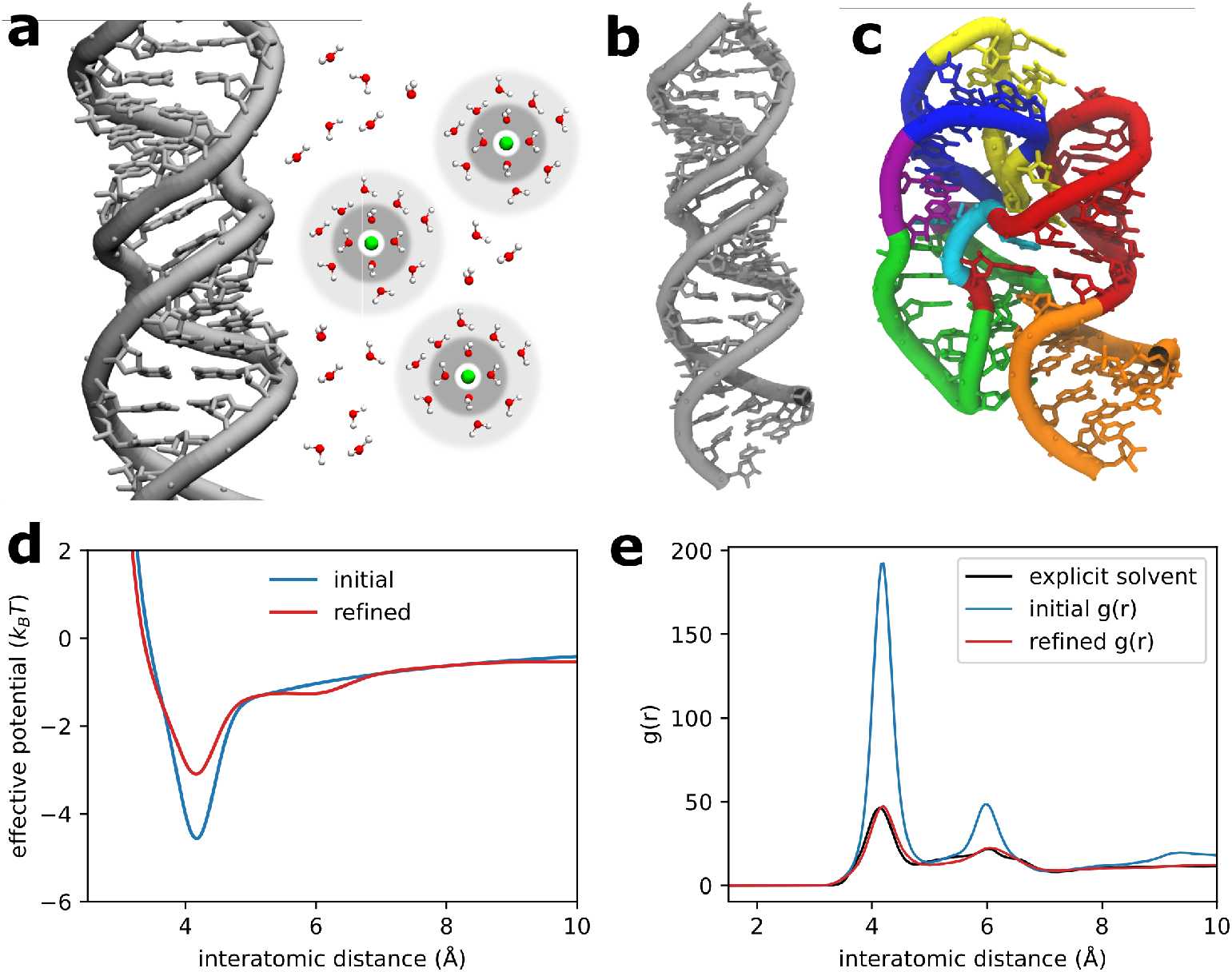
Describing the dynamics of diffuse ions: The SMOG-ion model. To study the influence of diffuse ions on large-scale molecular assemblies, we developed an all-atom model with simplified energetics (SMOG-ion) in which monovalent and divalent ions are explicitly represented. In this model, intramolecular interactions are defined by a structurebased model,^63,64^ partial charges are assigned to each atom, and effective potentials are introduced to account for the effects of ion-ion correlations and hydration. a) While chelated ions form strong interactions with biomolecules, diffuse ions (green beads) maintain hydration shells (gray rings) that prevent tight binding. b) Structure of helix 44 (h44) of the ribosomal small subunit, which was used as a test system for initial parameterization of the model. c) An rRNA 58-mer that was used for comparison and calibration against experimental measurements. d) Potential for diffuse Mg^2+^ interactions with highly-charge RNA oxygen atoms before (s0 parameters, blue) and after (s1 parameters, red) refinement against explicitsolvent simulations of h44. After refinement, the corresponding radial distribution function (panel e) agrees well with that obtained using the explicit-solvent model, which ensures the ionic shells are consistently described. For a list of modeled interactions, see Tabs. S1-S5. For comparison of *g*(*r*) for all interaction types, see Fig. S5. After comparison with explicitsolvent simulations, minor adjustments to Mg^2+^ and K^+^ interactions were subsequently introduced based on comparison with experiments for the 58-mer.^72^

In the second stage of refinement, a subset of the ion parameters was adjusted based on comparison with experimental measures of diffuse ionic distributions. For this, the preferential interaction coefficient of Mg^2+^ ions (Γ_2+_) was calculated from simulations of a 58nucleotide rRNA fragment^76^ (Fig. 1c and Fig. S2a). For consistency with the experimental conditions,^72^ we simulated the 58-mer with [MgCl_2_] = 1 mM and [KCl] = 150 mM. Even though the s1 model parameters recapitulate the ionic distributions predicted by the explicit-solvent model (Fig. 1e), Γ_2+_ was significantly underestimated for the 58-mer. Specifically, the predicted value of Γ_2+_ was 2.2 *±* 0.6, whereas the experimental value was 10.4.^72^

Since the underestimation of Γ_2+_ indicated an imbalance between K^+^ and Mg^2+^ association strengths with RNA,^3^ we introduced minor changes to the Mg^2+^ and K^+^ interactions with highly-electronegative RNA atoms. In explicit-solvent simulations of RNA, excess K^+^ ions are frequently partially dehydrated, which can artificially amplify the effective strength of K^+^-RNA interactions.^77,78^ This observation is at odds with NMR studies that have reported most K^+^ ions tend to remain fully hydrated. ^17^ Since the s1 parameter set was based on comparison with an explicit-solvent model, we removed the effective potential that stabilizes short-range (first outer shell) interactions between diffuse K^+^ ions and electronegative (*q < −*0.5) O and N atoms (yielding the s2 parameter set). While this increased

Γ_2+_ from 2.2 (s1) to 6.4 (s2), the persistent underestimation of Γ_2+_ suggested the effective potential for Mg^2+^ was also insufficiently stabilizing. Since Γ_2+_ is predominantly influenced by the distribution of diffuse Mg^2+^ ions around highly electronegative atoms,^39,77^ we introduced a small increase to the stability of these interactions in our model. Using estimates obtained from an energetic-based reweighting strategy,^ii^ we tested the effect of increasing the short-range (outer shell) Mg^2+^ interaction by 0.16 (s3 parameter set)^iii^. With this change, Γ_2+_ = 9.5 *±* 0.7, which is comparable to the experimental value of 10.4. Due to the uncertainty in the calculated Γ_2+_ values (see Methods), we decided to terminate the refinement process upon reaching this level of agreement. For all subsequent analysis, we used the s3 parameter set, which will simply be referred to as the SMOG-ion model.

### Spatial partitioning of diffuse ionic species

Our simplified model predicts condensation and spatial partitioning of ionic species, which is consistent with monovalent and divalent ions contributing differentially to the stability of secondary and tertiary structure in RNA. The predicted spatial distributions (Fig. 2) for the 58-mer are generally corroborated by prior crystallographic analysis and explicitsolvent simulations. In terms of biomolecular structure, K^+^ ions primarily populate the major grooves of the RNA, while Mg^2+^ ions appear to bridge interactions associated with tertiary structure.

**Figure 2:**
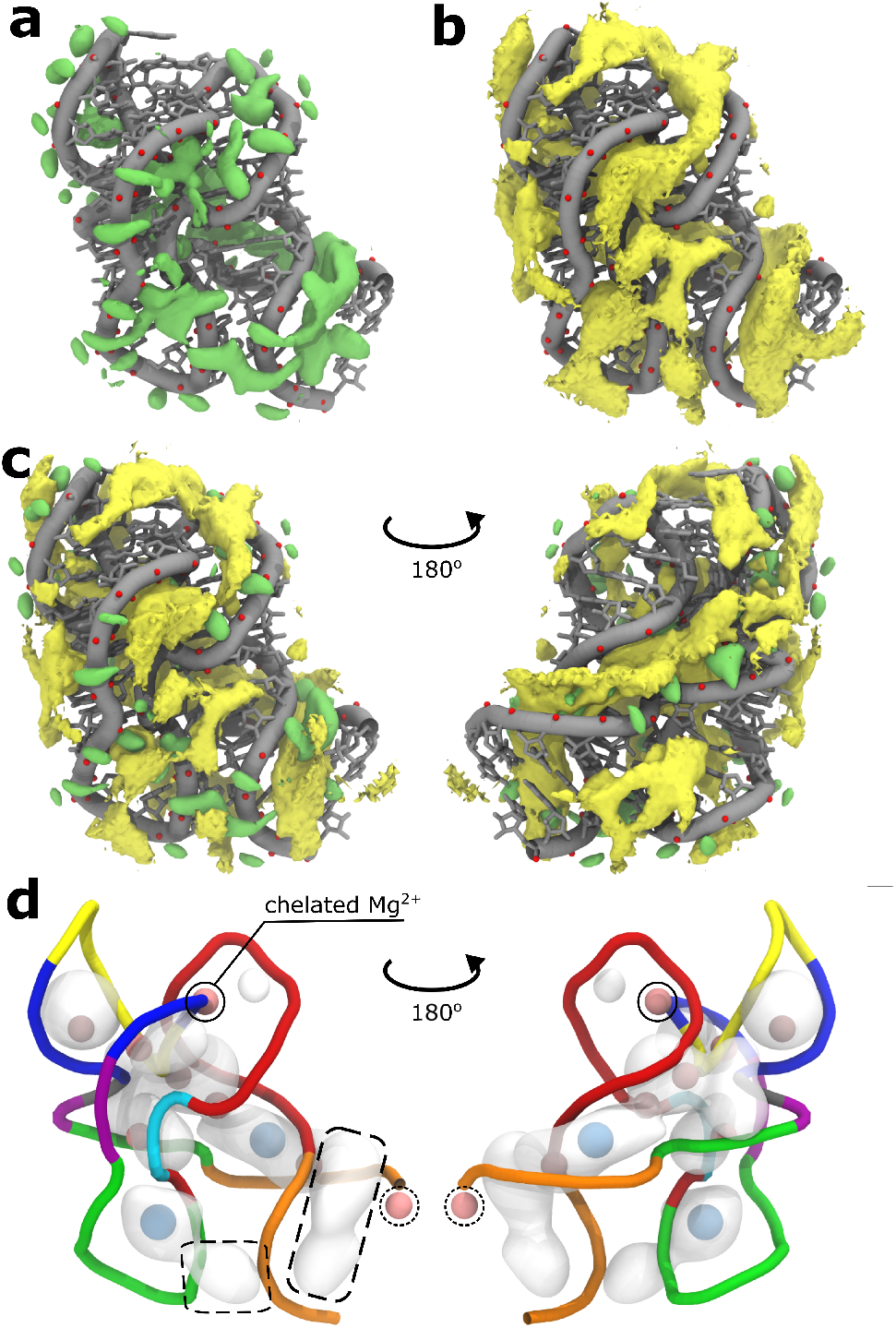
Characteristics of monovalent and divalent ion-RNA association. a) Spatial distribution function (SDF) of diffuse Mg^2+^ ions for the 58-mer ([MgCl_2_]=1 mM, [KCl]=150 mM). The isosurface represents a 1.3M concentration of Mg^2+^ (1300-fold enrichment over bulk). b) SDF for diffuse K^+^ ions (isosurface at 1.3M). c) The difference between the SDFs : 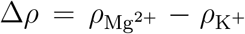 where *ρ*_i_ is the SDF of ion type i. The green isosurface shows preferential association of Mg^2+^ ions (Δ*ρ* = 1.3M), while yellow shows a preference for K^+^ ions (Δ*ρ* = 1.3M). K^+^ tends to populate the RNA grooves, consistent with its significant influence on the stability of secondary structure.^1^ Mg^2+^ ions are dominant along the RNA backbone, consistent with their contribution to tertiary structure formation.^2,3,26,27^ d) SMOG-ion model predicts population of crystallographically-reported ionic densities. SDF calculated for diffuse Mg^2+^ ions (chelated ion not included) after applying a Gaussian filter. Isosurface shown for Mg^2+^ concentration of 0.5M. Crystallographically assigned non-chelated Mg^2+^ ions (pink) and Os^3+^ ions (blue) are within the predicted regions of high diffuse Mg^2+^ densities, except a solitary Mg^2+^ ion near the terminal tail (dashed circle). In experiments, crystallographic contacts with the tail likely facilitate ion localization. In addition to predicting the crystallographic ions, there are two additional regions of high density (dashed boxes), which may further contribute to tertiary structure formation.

Before comparing the predicted ionic distributions with crystallographic data, it is necessary to describe the experimental assignment of ions. The crystal structure of the 58-mer^76^ (PDB ID: 1HC8) contains two asymmetric protein-RNA assemblies in the unit cell, and there are eight Mg^2+^ assignments that are common (Fig. 2d, pink beads)^iv^. Of these common assignments, Poisson–Boltzmann calculations indicate one is likely to be bound (Fig. 2d, circled. binding free energy Δ*G*_bind_ = *−*4.8 kcal/mol, Ref.^79^). Based on this, the system is typically described as possessing a single chelated Mg^2+^ ion.^76^ Since our model was parameterized for the study of diffuse ions, harmonic interactions were introduced to maintain the position of this single chelated Mg^2+^ ion. There is also a chelated K^+^ in^76^ that we restrain to its binding site. Finally, the crystal structure has two non-chelated Os^3+^ ions that were not included in our simulations.

The SMOG-ion model predicts high local concentrations of Mg^2+^ ions that coincide with the crystallographically-resolved positions of non-chelated multivalent ions. Specifically, the model predicts regions of high Mg^2+^ density that overlap with six of the seven non-chelated Mg^2+^ positions (Fig. 2). The only outlier is positioned near the terminal guanosine-5’- triphosphate residue (Fig. 2d, dashed circle). In the simulations, mobility of the tail impedes ion association. In contrast, in the experimental structure, crystallographic contacts with the tail likely reduce the flexibility, which in turn can facilitate a higher ion density. The model also predicts high Mg^2+^ densities that overlap with the experimentally assigned positions of both Os^3+^ ions (Fig. 2d, blue beads). Overall, consistency between the predicted SDFs and crystallographic analysis suggests that our model provides an accurate description of the local ionic environment.

The SMOG-ion model implicates differential contributions of monovalent and divalent ions to the stability of RNA. In our model, diffuse K^+^ and Mg^2+^ ions both populate the major groove of double-stranded RNA helices (Fig. 2a,b), as predicted by explicit-solvent simulations.^80,81^ This shows how both ionic species can contribute to the stability of secondary structure in RNA, a property also predicted by coarse-grained models.^39^ Consistent with explicit-solvent simulations,^77^ we also find that Mg^2+^ ions accumulate around each backbone phosphate group (Fig. 2c, green). As demonstrated in simulations with coarsegrained models,^26,27,39^ the observed distributions reinforce the notion that diffuse Mg^2+^ ions are a primary contributor to tertiary structure formation in RNA. Interestingly, the model predicts two regions of high Mg^2+^ density that do not coincide with assigned ion positions (Fig. 2d, boxed regions). Both of these regions span RNA segments that are distant in sequence. These high-density regions further suggest how Mg^2+^ ions may facilitate higherorder structure formation in RNA.

### Model captures concentration-dependent ionic atmosphere

Before applying the SMOG-ion model to a large biomolecular assembly, we evaluated its transferability by comparing the concentration dependence of Γ_2+_ with experimental values and values recently obtained with a coarse-grained model.^39^ For this, we considered two different RNA molecules: the 58-mer rRNA fragment^72^ and an adenine riboswitch.^73^ Comparing concentration-dependent values of Γ_2+_ allows one to ask whether the modeled parameters appropriately describe the competition of ionic species. It is important to note that the ion-RNA interaction strengths were assigned based on comparison with explicit-solvent simulation and a single experimental value of Γ_2+_. Accordingly, calculating Γ_2+_ values for multiple systems over a range of ion concentrations represents a blind test of the transferability of the model.

We first compared with previously-predicted and experimental^72^ values of Γ_2+_ for the 58-mer rRNA (Fig. 3a,b). While the SMOG-ion model was calibrated using the 58-mer with [MgCl_2_] = 1 mM and [KCl] = 150 mM, we find it accurately predicts the change in Γ_2+_ as a function of [MgCl_2_]. Over the studied concentrations (*∼*0.1-1mM), the experimental Γ_2+_ values change by 8.0, whereas the model predicts a change of 6.6. There is a systematic underestimation (*∼* 1) of the Γ_2+_ values at higher ion concentrations, though the statistical uncertainty in the theoretical values is comparable to this difference (Fig. 3b). In addition, the values of Γ_2+_ predicted by the SMOG-ion model agree well with those obtained with a leading coarse-grained model^39^ (Fig. 3b, purple dots). It is important to note that this analysis was performed for a hyperstable variant of the rRNA (U1061A), and comparisons are limited to higher experimental ionic concentrations. Since this RNA is known to remain folded under these conditions, this set of comparisons allows us to specifically assess the accuracy of the ion parameters, in a manner that is independent of the biomolecular potential. However, it is possible that the experimental values also reflect concentration-dependent effects on RNA structure that are not addressed here. With these experimental and statistical uncertainties in mind, one may consider the residual differences between Γ_2+_ values to be minor.

**Figure 3:**
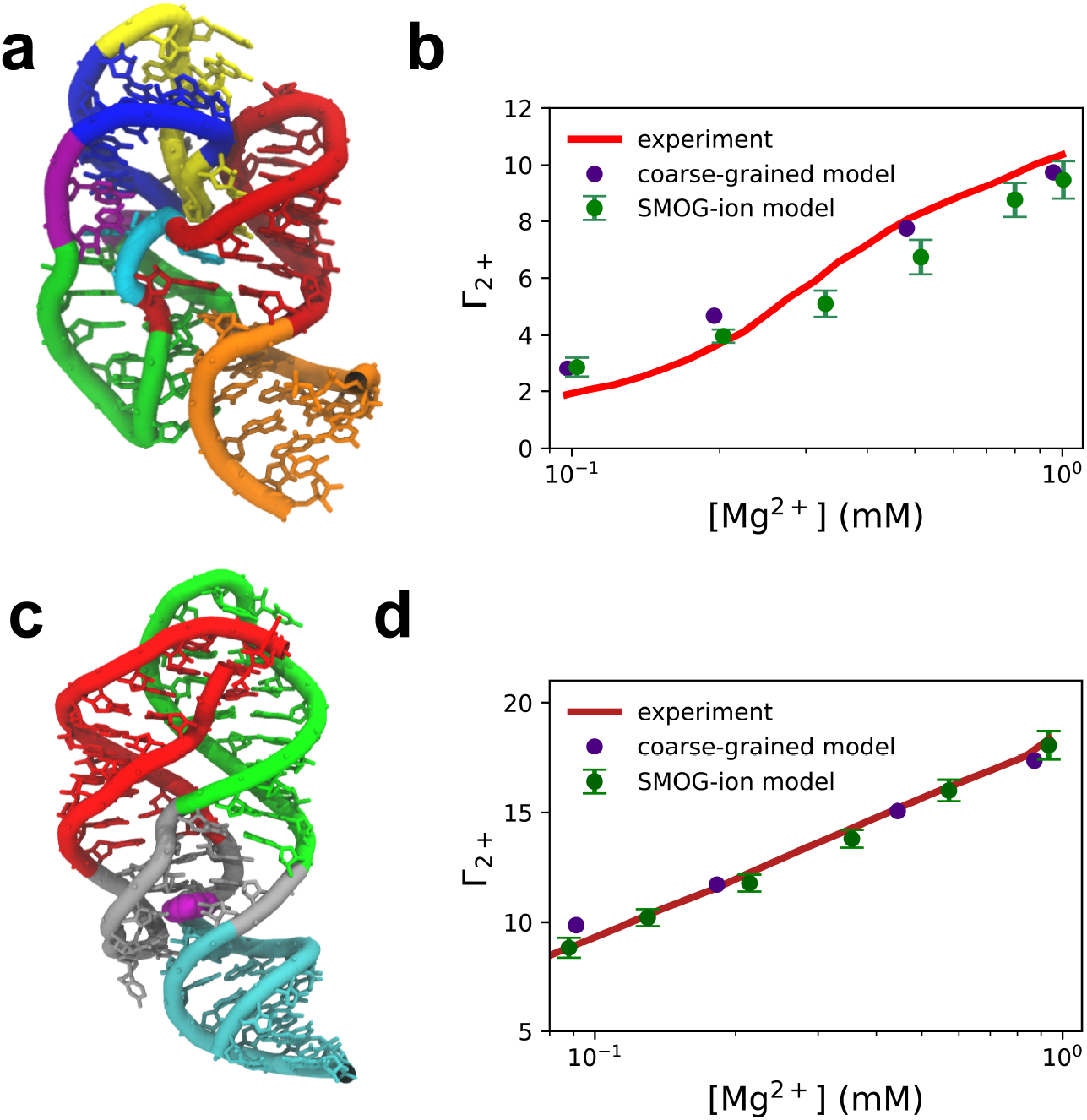
Simplified model captures concentration-dependent ion association. a) Tertiary structure of the 58-mer rRNA fragment^76^ (colored as in Fig. 1c). b) Preferential interaction coefficient (Γ_2+_) for the 58-mer ([KCl] = 150mM), shown for the the SMOG-ion model, previous experimental values^72^ and predictions from a coarse-grained model of Nguyen et el.^39^ While our model was parameterized based on comparison with the experimental value of Γ_2+_ at [MgCl_2_] = 1mM, the predicted concentration dependence (green dots) follows the experimental behavior (red curve) and agrees well with previous predictions from the coarse-grained model^39^ (purple dots). c) Tertiary structure of the adenine riboswitch,^73^ colored as in Fig. S2b. d) The value of Γ_2+_ for the adenine riboswitch ([KCl] = 50mM), obtained with the SMOG-ion model (green dots), experimental measurements^73^ (dark red curve) and the coarse-grained model of Nguyen et al.^39^ (purple dots). There is excellent agreement between the values, even though the riboswitch was not used for model parameterization, and our model parameters were established using benchmark systems at higher value of [KCl] (100-150 vs. 50 mM). Accordingly, these comparisons support the transferrability of the model to other RNA systems and ionic concentrations. Error bars in (b) and (d) represent the standard deviation of Γ_2+_ calculated from 20 replicate simulations.

Applying the model to the adenine riboswitch^v^ (Fig. 3c) demonstrates the transferability of the ion parameters. Since the riboswitch was not utilized for any aspect of parameter refinement, it serves as a blind test of the predictive capabilities of the model. In addition, the experiments were performed at lower values of [KCl] for the adenine riboswitch than forthe 58-mer (50 mM^73^ vs. 150 mM^72^). Accordingly, this comparison implicitly evaluates the predicted concentration-dependent influence of monovalent ions on Mg^2+^-RNA association. We find that the predicted Γ_2+_ values agree very well with the experimental measurements,^73^ where the level of agreement is comparable to predictions from the coarse-grained model of Nguyen et al.^39^(Fig. 3d). Combined with our analysis of the 58-mer, this demonstrates the ability of the SMOG-ion model to accurately estimate the energetics and association of diffuse ions.

### Diffuse ions control tRNA kinetics on the ribosome

After benchmarking the SMOG-ion model, we used it to investigate how diffuse ions contribute to a large-scale conformational transitions in the ribosome: aminoacyl-tRNA (aa-tRNA) accommodation. After delivery of aa-tRNA to the ribosome by EF-Tu, the accommodation process allows the tRNA to fully bind the ribosome, where this step is responsible for proofreading the incoming tRNA molecule.^82^ Here, we simulated the first step of aa-tRNA accommodation, called elbow accommodation (Fig. 4A). This large-scale (*∼* 20 *−* 30Å) conformational rearrangement has been extensively studied using electrostatics-free models^83–86^ and implicit-ion models.^87^ Explicit-solvent simulations have also been used to calculate diffusion coefficients^88,89^ and to perform nanosecond-scale targeted simulations of this process. ^90^ This body of work has shown that Helix 89 (H89) introduces a pronounced sterically-induced free-energy barrier^83–86^ that can be amplified by direct electrostatic interactions between aa-tRNA and H89.^87^ Electrostatic and solvent interactions between tRNA and H89 can also introduce energetic roughness that leads to coordinate-dependent diffusive properties of tRNA.^89^ There have also been experimental insights into the roles of ions. For example, anomalous scattering data has been used to identify the composition of bound ions on the ribosome^91^ and changes in solvent conditions have been shown to dramatically alter the kinetics of accommodation.^15,92^ Here, the SMOG-ion model provides complementary insights into tRNA dynamics by specifically isolating the influence of diffuse ions on the kinetics of accommodation.

**Figure 4:**
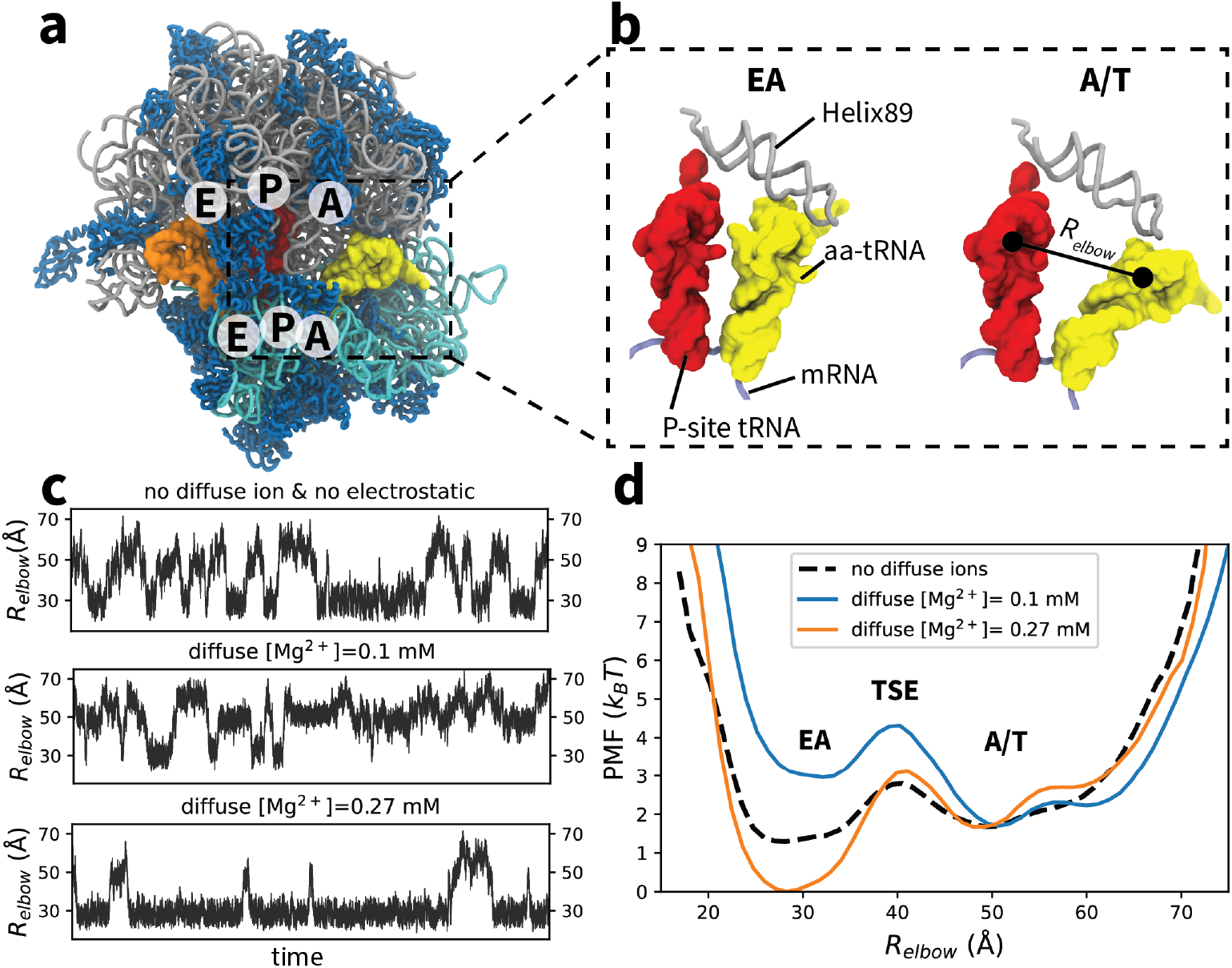
Diffuse ions facilitate aa-tRNA elbow accommodation. a) Structure of the 70S ribosome with 23S rRNA in gray, 16S rRNA in cyan and proteins in blue. The ribosome is shown in complex with A/T-configured aa-tRNA (yellow), P-site tRNA (red), Esite tRNA (orange) and mRNA (purple). b) aa-tRNA elbow accommodation involves a ~25Å rearrangement between A/T and EA configurations. *R*_elbow_ is the distance between the O3’ atoms of U60 in the aa-tRNA and U8 in the P-site tRNA.^84^ c) Representative trajectories of simulations using all-atom structure-based models with different treatment of electrostatics and diffuse ions. In all systems, spontaneous transitions are observed between the A/T and EA ensembles. The electrostatics-free and ion-free model^84^ (top) is used as a reference, where there is roughly equal probability of adopting either endpoint. Depending on the ionic concentration (middle, bottom), the range of accessible tRNA positions is shifted. d) The potential of mean force (PMF = *k*_B_*T* ln(*P*)) for each model. In the absence of electrostatics and diffuse ions (dashed line), the free energies of the A/T to EA ensembles are comparable. When explicit electrostatics is included, and [KCl] = 100mM with a concentration of 0.1 mM for diffuse Mg^2+^ ions (blue), the landscape is shifted towards the A/T ensemble. At higher concentrations of diffuse Mg^2+^ ions (orange curve and Fig. S7) the free-energy of the transition state ensemble (TSE) is reduced and the landscape is shifted towards the EA ensemble, which corresponds to aa-tRNA binding of the ribosome.

To examine the role of diffuse ions during aa-tRNA elbow accommodation, we performed simulation of the complete 70S ribosome using the SMOG-ion model, and compared the dynamics for different ionic strengths. Consistent with earlier analysis,^84^ we calculated the free-energy barrier as a function of *R*_elbow_: the distance between the O3’ atoms of U60 of aa-tRNA and U8 of P-site tRNA. In these simulations, spontaneous transitions between the A/T (*R*_elbow_*∼* 50-60Å) and EA ensembles (*R*_elbow_*∼* 30Å) are observed (with ions present, or absent). When the electrostatics-free model is used (labeled “SMOG model” in Tab. S5), the free energy of the A/T and EA ensembles is comparable (Fig. 4d, black). When electrostatics and monovalent ions ([KCl] = 100mM) are included, along with bound Mg^2+^ ions (no diffuse Mg^2+^), the energy landscape is shifted towards the A/T ensemble, and there are only transient excursions in the direction of the EA ensemble (Fig. S6). This shows how, even when monovalent ion screening is accounted for, there is strong electrostatic repulsions between the aa-tRNA and the ribosome. When the bulk concentration of diffuse Mg^2+^ ions is only 0.1 mM, aa-tRNA is found to stably populate both the A/T and EA ensembles, where the free-energy barrier between A/T to EA is *∼* 3*k*_*B*_*T* (Fig. 4d, blue). Interestingly, this is slightly larger than the barrier obtained with the electrostatic-free model (*∼* 1.5*k*_*B*_*T*), illustrating the residual RNA-RNA repulsion that is present under these conditions. When the bulk concentration of Mg^2+^ ions is increased to 0.27 mM, the free-energy barrier is reduced by *∼* 1.5 *k*_*B*_*T* (Fig. 4d, orange), consistent with further attenuation of electrostatic repulsions between H89 and aa-tRNA. The EA ensemble is also stabilized at this higher concentration of Mg^2+^ (Fig. 4d, orange). When the concentration of diffuse Mg^2+^ ions is further increased, aa-tRNA is observed to strongly favor the EA ensemble over A/T (Fig. S7), where the dynamics is suggestive of a downhill energy landscape. Together, these results illustrate the strong dependence of aa-tRNA kinetics and thermodynamics on the precise concentration of diffuse Mg^2+^ ions.

While the current analysis reveals a clear energetic role of diffuse ions, these observa-tions suggest many interesting avenues for continued investigation. For example, similar to a coarse-grained model,^39^ one may extend the SMOG-ion model to describe binding of chelated ion, which would be necessary to account for their effects on RNA stability. In this regard, while the flexibility of the ribosome is well described by the SMOG-ion model (Fig. S8), the overall stability will need further characterization and parameterization. That is, it will be important to better understand the scale of stabilization imparted by the combination of structure-based energetics and non-specific electrostatic interactions. Resolving this limitation can also open the possibility of identifying localized disorder events, which would have the potential to influence substeps of elongation. Another feature of the present model is that it was tuned so that the A/T and EA ensembles are of comparable free-energy in the absence of electrostatics. This strategy was applied, in order for the ion-free simulations to provide a reference distribution, against which the perturbative effects of ionic concentration changes could be compared. In future models, one may envision relaxing the explicit stability of the accommodated (EA) basin. Such a change would shift the landscape towards the A/T ensemble, where higher ionic concentrations (i.e. closer to biological values) will likely be necessary for favorable A-site binding to occur. As a final example, it will be interesting to characterize the detailed influence of Elongation Factor-Tu when diffuse ions are present. While screened-electrostatic models have shown that EF-Tu can facilitate the accommodation process through steric effects,^87^ it is possible that diffuse ions will further modulate this influence on the incoming tRNA molecules.

### How ion-mediated interactions regulate tRNA-ribosome dynamics

To provide structural insights into the mechanisms by which diffuse ions can alter the energy landscape of the ribosome, we analyzed the statistical properties of ion-mediated interactions. Specifically, we identified all Mg^2+^-mediated interactions that are formed between highly negatively charged atoms of the aa-tRNA and rRNA of the large subunit (LSU). Here, an interaction is defined as “ion-mediated” if an Mg^2+^ ion is simultaneously within 5 Å of the aa-tRNA and LSU. To connect this analysis with the observed changes in the free-energy landscape (Fig. 4d), we evaluated the average number of ion-mediated interactions formed in the EA, transition state (TSE) and A/T ensembles. This reveals several ion association “hot spots” on the ribosome that form significant numbers of ion-mediated contacts with the aa-tRNA molecule. We also find specific regions for which there are clear concentrationdependent interactions with tRNA, which provides a molecular explanation for the ion-dependent effects on accommodation kinetics.

The largest number of ion-mediated interactions are formed between the tRNA molecule and H89. As noted above, the excluded volume of H89 has been shown to introduce a sterically-induced free-energy barrier during elbow accommodation,^84–86^ where direct electrostatic interactions can amplify the barrier height.^87^ Here, we find distinct sets of ion-mediate interactions are transiently formed with H89 during the accommodation process (Fig. 5). At the early stages of accommodation, the aa-tRNA approaches the stem loop of H89 (C2471-C2474; orange in Fig. 5c). Upon reaching the TSE, the aa-tRNA forms up to around three ion-mediated interactions with individual residues (e.g. C2472), where the number of interactions increases with the concentration of divalent ions. This is consistent with the observation that the free-energy barrier is smaller (reduced by 1.5*k*_*B*_*T*) for the higher concentration of diffuse ions (Fig. 4d; blue vs. orange curves). After the tRNA overcomes the free-energy barrier and enters the EA ensemble, these transient ion-mediated interactions dissolve, which is expected due to the increased distance between the stem loop and the tRNA molecule.

**Figure 5:**
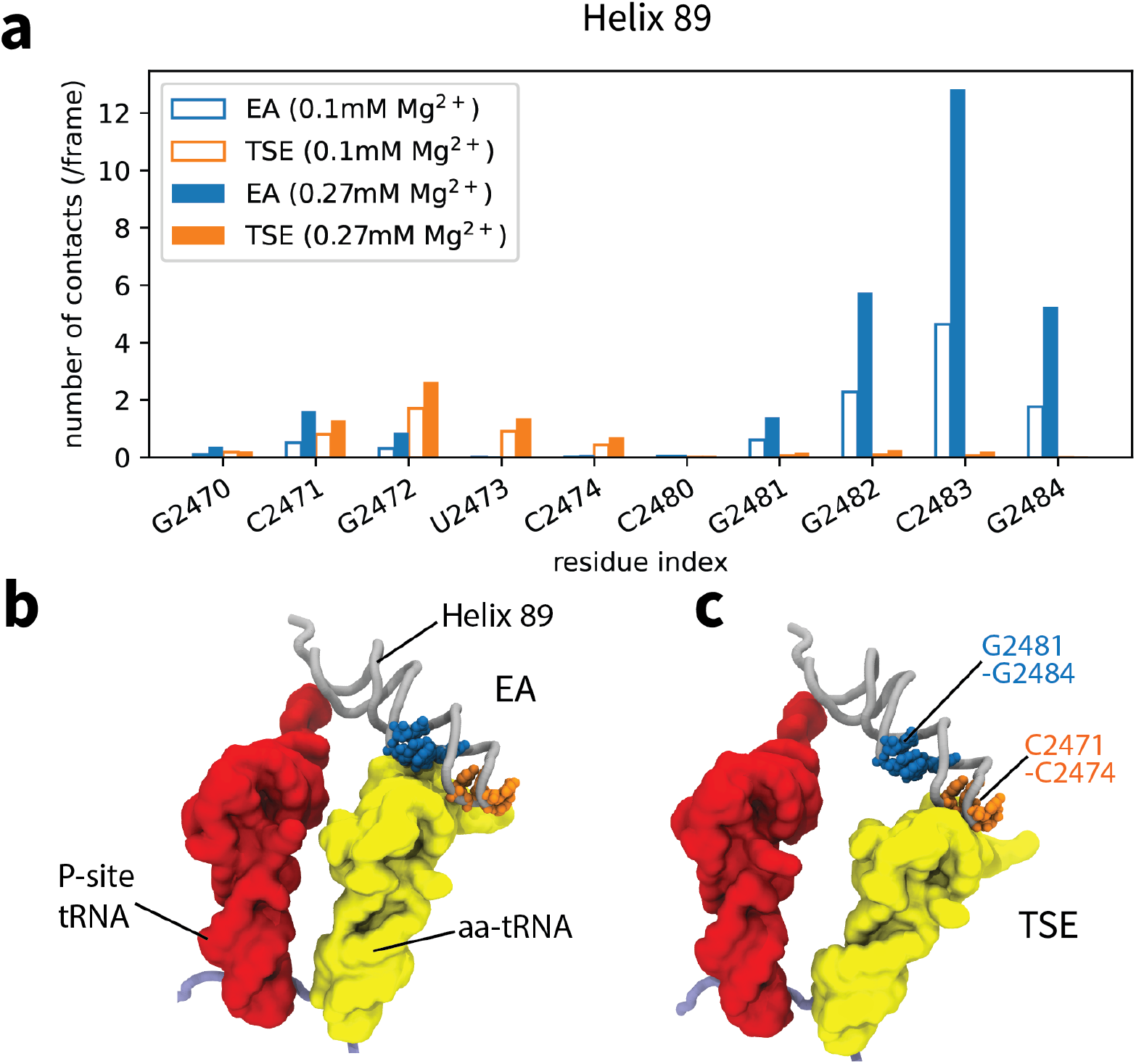
Mg^2+^-mediated interactions explain changes in the energy landscape. a) The average number of Mg^2+^-mediated interactions between aa-tRNA and H89, calculated for the EA ensemble (blue) and Transition State Ensemble (TSE: orange). Solid bars correspond to a diffuse Mg^2+^ concentration of 0.27 mM, while empty bars correspond to 0.1 mM. In the TSE (panel c), transient ion-mediated interactions are formed between the aa-tRNA and the stem loop of H89 (C2471-C2474; orange residues). There is also a slight increase at higher concentrations of Mg^2+^, which is consistent with the smaller free-energy barrier (Fig. 4). Upon reaching the EA ensemble (panel b), ion-mediate interactions with the stem loop dissolve, while a larger number of ion-mediated interactions is then formed with the base of H89 (G2481-G2484; blue residues). There is also a sharp increase in the number of ion-mediated interactions at higher concentrations, consistent with the larger energetic effect of ions on the free energy of the EA (ribosome-bound) ensemble.

As the tRNA molecule enters the EA ensemble, a second set of ion-mediated interactions is formed. Specifically, there is a large increase in the number of interactions with the base of H89 (Fig. 5b; G2481-G2484 in blue). A small number of contacts is also formed with H92 (Fig. S9). Together, there are more ion-mediate interactions formed in the EA ensemble than in the TSE. The number of contacts formed in the EA ensemble also exhibits a stronger concentration dependence, where an individual residue (C2483) gains up to eight additional ion-mediated interactions at higher concentrations. This stark concentration dependence of ion-mediated contacts rationalizes the more pronounced increase in stability of the EA ensemble (*∼* 3*k*_*B*_*T*), relative to changes found for the TSE (*∼* 1.5*k*_*B*_*T*).

In addition to the influence of H89, our simulations also implicate a minor influence of the L11 stalk on accommodation kinetics. The L11 stalk is a flexible and extended region that is located at the periphery of the ribosome (Fig. S10), and it is essential for the binding of elongation factors (EF-Tu and EF-G).^93–96^ It is also the binding site of thiostrepton, which inhibits translocation by preventing phosphate release. ^97,98^ Previous simulations have shown that changes in L11 flexibility can impact the free-energy barrier associated with the elbow accommodation,^85^ where L11 is able to confine the aa-tRNA motion within the A/T ensemble. To complement this effect, we find that Mg^2+^-mediated interactions between tRNA and L11 are more common in the A/T ensemble and TSE (Fig. S11). In the EA ensemble, ion-mediated interactions with L11 are rare, due to the long distance between these structural elements. At higher concentrations, we find a modest increase in the number of ion-mediated interactions in the TSE (*∼* 0.3) and a marginal decrease in the A/T ensemble (*∼* 0.15). Together, these small changes may further contribute to the apparent reduction of the free-energy barrier at higher Mg^2+^ concentrations (Fig. 4d).

## Discussion

### Strategies for studying diffuse ions

The accuracy of ionic models is often evaluated based on studies of the adenine riboswitch and 58-mer rRNA. The reason for this choice is that high-quality experimental measures are available that describe the distribution of monovalent (K^+^ and Cl^*−*^) and divalent (Mg^2+^) ions around these RNA molecules.^72,73^ As described in the results, even though parameterization was based on experimental comparisons for a single ionic concentration, we find that the SMOG-ion model is able to capture the experimentally-measured concentration dependence of Γ_2+_ for both systems. In terms of these common benchmarks, the SMOG-ion model provides a level of agreement that is comparable to other available models. Lammert *et al*. found that explicit-solvent simulations of the 58-mer^81^ overestimated Γ_2+_ by 2-3. However, due to the small system size in that study, there was a significant uncertainty in the calculated bulk ion concentration. In terms of coarse-grained approaches, Nguyen *et al*.^39^ used theory based on the reference interaction site model to develop effective potentials for divalent ions. This representation was able to predict Γ_2+_ to within a value of 1 for both systems (Fig. 3), which is comparable to the accuracy of SMOG-ion. While their use of pair-wise effective potentials is similar to our approach, that model employed an implicit representation of monovalent ions. As a result, it was not tractable to calculate Γ_2+_ for the lowest concentrations of Mg^2+^ reported by Nguyen et al, since this would necessitate the use of a prohibitively large system of monovalent ions. Γ_2+_ values have also been accurately predicted using a generalized Manning condensation model proposed by Hayes *et al*.^99^ Overall, we find that the SMOG-ion model is able to predict Γ_2+_ with accuracy that is comparable to leading theoretical models.

While numerous models can accurately predict experimental Γ_2+_ values, each is suited to address specific physical questions. In coarse-grained models developed by Thirumalai and colleagues,^26,27,39,42,54^ a major focus has been to predict ion-dependent folding dynamics of large RNA molecules. Since folding can be described well with coarse-grained representations,^100^ three-site models have been appropriate for those purposes. In the study of Nguyen et al.,^39^ an objective was to understand the dynamics of chelated ions during folding, which necessitated the proper treatment of inner-shell interactions with Mg^2+^. In the study of Lammert *et al*.,^81^ explicit-solvent simulations were used to identify how small-scale (short time) structural fluctuations were correlated with individual ion association events. With this finely-detailed question in mind, a higher-resolution model was warranted. In contrast, Hayes *et al*.^99^ focused on more general questions pertaining to the utility of Manning theory to quantitatively predict ion condensation in asymmetric molecular systems, where an intermediate-resolution model is suitable.

Here, our aim is to study structural dynamics that involve transient formation of ion-mediated interactions. As an initial application, we have described how transient ion-association events are correlated with large-scale (*∼* 30 Å) rearrangements of tRNA in the ribosome. Since numerous studies have shown that detailed steric interactions strongly influence the dynamics of tRNA in the ribosome, it was necessary for the SMOG-ion model to employ an all-atom representation of the biomolecules, while coarse-graining over the solvent.

### Modeling the factors that control biomolecular assemblies

A major challenge in the study of ribonucleoprotein assemblies has been to precisely describe the influence of ions during large-scale conformational transitions. While there are efforts to refine ion parameters for use with explicit-solvent models, the large size of RNP assemblies (often MDa scales), combined with the slow timescales (*µ*s-ms-s), makes direct simulation of the dynamics intractable with conventional models. In an effort to bridge our understanding of large-scale dynamics and ionic effects, we propose the SMOG-ion model, which is far less computationally expensive than explicit-solvent simulations. Due to the simple functional form, these simulations scale to thousands of compute cores for large assemblies (Fig. S12), and they can be performed effectively using modern GPU resources.^101^ With these levels of performance, modest-sized compute clusters are sufficient to perform simulations that describe millisecond effective timescales for MDa-scale assemblies. Accordingly, it is now becoming feasible to explore the impact of ions on large-scale conformational rearrangements. For the SMOG-ion model, all-atom resolution was employed since previous studies have repeatedly shown how sterically-induced free-energy barriers can limit the kinetics of the ribosome. For example, multi-basin structure-based models have demonstrated how the steric composition of the A-site finger controls the scale of the free-energy barrier associated with A/P hybrid formation.^102^ During P/E hybrid formation, similar models have shown that the kinetics depends critically on the precise steric representation of the N-terminal tail of protein L33.^103^ In these examples, direct perturbations to the steric representation in the model revealed the central influence of excluded volume. Since the contribution of sterics will be robust, so long as atomic resolution is included, it is expected that the positions of these barriers will also be robust to the energetic details of the model. However, by applying more energetically-complete models, such as the SMOG-ion model, future studies will be able to further delineate the factors that control the scale of each barrier, and therefore the biological kinetics.

To complement the insights provided by the SMOG-ion model, explicit-solvent simulations and quantum mechanical methods can provide highly-detailed descriptions of biomolecular interactions within larger assemblies. For the ribosome, explicit-solvent models have been widely used to quantify energetics of bimolecular binding events,^104^ or small-scale (e.g. single-residue) structural rearrangements.^105^ In other studies, they have been used to quantify the stabilizing interactions between different components of the ribosome, which have included studies of the L1 stalk,^106,107^ interactions between the 3’-CCA tail of tRNA and its binding sites,^108^ as well as between tRNA and the ribosome,^109^ synthetases^110^ and elongation factors.^111^ At an even higher level of resolution, quantum mechanical techniques are available to study chemical reactions.^112,113^

The stark differences between simplified/coarse-grained models and explicit-solvent techniques often makes it difficult to integrate the results within a common quantitative framework. Here, we present a model that is substantially more energetically detailed than traditional structure-based models, while also being far simpler than explicit-solvent models. We anticipate that these intermediate-resolution models will help establish a more comprehensive understanding of RNP dynamics that will bridge the gap between detailed insights arising from explicit-solvent simulations and larger-scale processes that can be described by coarse-grained techniques. For example, one may use high-resolution methods to quantify precise energetic features (e.g. binding energetics, pH effects, etc) and then encode these features into a SMOG-ion model variant. Through this, it would be possible to identify the effects of these localized interactions on larger-scale motions. Thus, building on the present results, one may methodically construct a comprehensive physical-chemical model that bridges these disparate length and time scales.

## Conclusions

Attributing specific roles to diffuse ions in ribonucleoprotein assemblies has remained elusive. While there has been notable progress in the study of monomeric systems and RNA folding,^27^ unambiguously identifying specific physical-chemical effects of diffuse ions in assemblies continues to be challenging. As a step in this direction, we developed and employed a model that provides an explicit treatment of non-Hydrogen atoms and ions, while providing an implicit treatment of solvent. We find that our model is able to capture experimental measures of the ionic environment for prototypical RNA systems, which motivated our application to a more complex system: the ribosome. By comparing the dynamics with a range of theoretical representations, we identify how diffuse-ion-mediated interactions can coordinate a large-scale rearrangement in the ribosome. With this foundation, the study provides a framework for identifying the ways in which diffuse ions help regulate the dynamics of complex biomolecular assemblies.

## Acknowledgement

PCW was supported by NSF grant MCB-1915843. Work at the Center for Theoretical Biological Physics was also supported by the NSF (Grant PHY-2019745). We acknowledge generous support provided Northeastern University Discovery Cluster, the C3DDB cluster. We also acknowledge support from the AMD HPC Fund. We thank Yang Wang for comments and proofreading the manuscript.

## Supporting Information Available

The following files are available free of charge.

• SI.pdf: Additional computational details, methods and figures.

## Notes

The authors declare no competing financial interest.

## Footnotes

i With the large dimensions of the simulated systems, the free volume (i.e. volume excluding the RNA) only differs from the total volume by less than 0.01%.

ii Γ_2+_ was estimated for alternate parameter values by weighting each simulated frame by exp(*β*Δ*E*),where Δ*E* is the change in system energy upon introduction of a modified parameter.

iii The reduced energy unit *E* is equal to 2*k*_B_*T*. For Mg^2+^-O interactions, *B*_1_ was changed from -0.73 to -0.89. For Mg^2+^-N interactions, it was changed from -0.86 to -1.02.

iv 21 coordinates of Mg^2+^ ions were reported in the two asymmetric assemblies of the 58-mer (chain C and chain D in PDB 1HC8). Alignment of the monomers reveals 13 distinct binding positions, where 8 are occupied in both RNA molecules.

v with ligand bound

## References

(1) Pyle, A. M. Metal ions in the structure and function of RNA. J. Biol. Inorg. Chem. 2002, 7, 679–690.

(2) Woodson, S. A. Metal ions and RNA folding: a highly charged topic with a dynamic future. Curr. Opin. Chem. Biol. 2005, 9, 104–109.

(3) Draper, D. E. A guide to ions and RNA structure. RNA 2004, 10, 335–343.

(4) Draper, D. E.; Grilley, D.; Soto, A. M. Ions and RNA folding. Annu. Rev. Biophys. Biomol. Struct. 2005, 34, 221–243.

(5) Schroeder, R.; Barta, A.; Semrad, K. Strategies for RNA folding and assembly. Nat. Rev. Mol. Cell Biol. 2004, 5, 908–919.

(6) Wan, Y.; Qu, K.; Ouyang, Z.; Kertesz, M.; Li, J.; Tibshirani, R.; Makino, D. L.; Nutter, R. C.; Segal, E.; Chang, H. Y. Genome-wide measurement of RNA folding energies. Mol. Cell 2012, 48, 169–181.

(7) Izatt, R. M.; Christensen, J. J.; Rytting, J. H. Sites and thermodynamic quantities associated with proton and metal ion interaction with ribonucleic acid, deoxyribonucleic acid, and their constituent bases, nucleosides, and nucleotides. Chem. Rev. 1971, 71, 439–481.

(8) Shamsi, M. H.; Kraatz, H. B. Interactions of metal ions with DNA and some applications. J. Inorg. Organomet. Polym. Mater. 2013, 23, 4–23.

(9) Abeysirigunawardena, S. C.; Woodson, S. A. Differential effects of ribosomal proteins and Mg^2+^ions on a conformational switch during 30S ribosome 5’-domain assembly. RNA 2015, 21, 1859–1865.

(10) Kim, H. D.; Nienhaus, G. U.; Ha, T.; Orr, J. W.; Williamson, J. R.; Chu, S. Mg^2+^-dependent conformational change of RNA studied by fluorescence correlation and FRET on immobilized single molecules. Proc. Natl. Acad. Sci. USA 2002, 99, 4284– 4289.

(11) Klein, D. J.; Moore, P. B.; Steitz, T. A. The contribution of metal ions to the structural stability of the large ribosomal subunit. RNA 2004, 10, 1366–1379.

(12) Gesteland, R. F. Unfolding of Escherichia coli ribosomes by removal of magnesium. J. Mol. Biol. 1966, 18, 356–371.

(13) Kim, H. D.; Puglisi, J. D.; Chu, S. Fluctuations of transfer RNAs between classical and hybrid states. Biophys. J. 2007, 93, 3575–3582.

(14) Holtkamp, W.; Cunha, C. E.; Peske, F.; Konevega, A. L.; Wintermeyer, W.; Rodnina, M. V. GTP hydrolysis by EF-G synchronizes tRNA movement on small and large ribosomal subunits. EMBO J. 2014, 33, 1073–1085.

(15) Wohlgemuth, I.; Pohl, C.; Rodnina, M. V. Optimization of speed and accuracy of decoding in translation. EMBO J. 2010, 29, 3701–3709.

(16) Johansson, M.; Zhang, J.; Ehrenberg, M. Genetic code translation displays a linear trade-off between efficiency and accuracy of tRNA selection. Proc. Natl. Acad. Sci. USA 2012, 109, 131–136.

(17) Braunlin, W. H. NMR studies of cation-binding environments on nucleic acids. Adv. Biophys. Chem. 1995, 5, 89–140.

(18) Reid, S. S.; Cowan, J. A. Biostructural Chemistry of Magnesium Ion: Characterization of the Weak Binding Sites on tRNAPhe(yeast). Implications for Conformational Change and Activity. Biochemistry 1990, 29, 6025–6032.

(19) Cowan, J. A. Coordination chemistry of Mg^2+^ and 5S rRNA (Escherichia coli): Binding parameters, ligand symmetry, and implications for activity. J. Am. Chem. Soc. 1991, 113, 675–676.

(20) Kowerko, D.; König, S. L.; Skilandat, M.; Kruschel, D.; Hadzic, M. C.; Cardo, L.; Sigel, R. K. Cation-induced kinetic heterogeneity of the intron-exon recognition in single group II introns. Proc. Natl. Acad. Sci. U. S. A. 2015, 112, 3403–3408.

(21) Caminiti, R.; Licheri, G.; Piccaluga, G.; Pinna, G. X-ray diffraction study of MgCl_2_ aqueous solutions. J. Appl. Crystallogr. 1979, 12, 34–38.

(22) Weingärtner, H.; Müller, K. J.; Hertz, H. G.; Edge, A. V.; Mills, R. Unusual behavior of transport coefficients in aqueous solutions of zinc chloride at 25°C. J. Phys. Chem. 1984, 88, 2173–2178.

(23) Bleuzen, A.; Pittet, P. A.; Helm, L.; Merbach, A. E. Water exchange on magnesium(II) in aqueous solution: A variable temperature and pressure ^17^O NMR study. Magn. Reson. Chem. 1997, 35, 765–773.

(24) Cole, P. E.; Yang, S. K.; Crothers, D. M. Conformational changes of transfer ribonucleic acid. Equilibrium phase diagrams. Biochemistry 1972, 11, 4358–4368.

(25) Urbanke, C.; Römer, R.; Maass, G. Tertiary structure of tRNA^Phe^(Yeast): Kinetics and electrostatic repulsion. Eur. J. Biochem. 1975, 55, 439–444.

(26) Denesyuk, N. A.; Thirumalai, D. How do metal ions direct ribozyme folding? Nat. Chem. 2015, 7, 793–801.

(27) Hori, N.; Denesyuk, N. A.; Thirumalai, D. Shape changes and cooperativity in the folding of the central domain of the 16S ribosomal RNA. Proc. Natl. Acad. Sci. USA 2021, 118, e2020837118.

(28) Fuoss, R. M.; Katchalsky, A.; Lifson, S. The potential of an infinite rod-like molecule and the distribution of the counter ions. Proc. Natl. Acad. Sci. USA 1951, 37, 579–589.

(29) Honig, B.; Nicholls, A. Classical electrostatics in biology and chemistry. Science 1995, 268, 1144–1149.

(30) Forsman, J. A simple correlation-corrected Poisson-Boltzmann theory. J. Phys. Chem. B 2004, 108, 9236–9245.

(31) Manning, G. S. The molecular theory of polyelectrolyte solutions with applications to the electrostatic properties of polynucleotides. Q. Rev. Biophys. 1978, 11, 179–246.

(32) Mohanty, U.; Ninham, B. W.; Oppenheim, I. Dressed polyions, counterion condensation, and adsorption excess in polyelectrolyte solutions. Proc. Natl. Acad. Sci. USA 1996, 93, 4342–4344.

(33) Manning, G. S.; Mohanty, U. Counterion condensation on ionic oligomers. Phys. A: Stat. Mech. and its Appl. 1997, 247, 196–204.

(34) Mohanty, U.; Spasic, A.; Kim, H. D.; Chu, S. Ion atmosphere of three-way junction nucleic acid. J. Phys. Chem. B 2005, 109, 21369–21374.

(35) Taubes, C. H.; Mohanty, U.; Chu, S. Ion atmosphere around nucleic acid. J. Phys. Chem. B 2005, 109, 21267–21272.

(36) Spasic, A.; Mohanty, U. Counterion condensation in nucleic acid. Adv. Chem. Phys. 2008, 139, 139–176.

(37) Hirata, F.; Rossky, P. J. An extended rism equation for molecular polar fluids. Chem. Phys. Lett. 1981, 83, 329–334.

(38) Pettitt, B. M.; Rossky, P. J. Integral equation predictions of liquid state structure for waterlike intermolecular potentials. J. Chem. Phys. 1982, 77, 1451–1457.

(39) Nguyen, H. T.; Hori, N.; Thirumalai, D. Theory and simulations for RNA folding in mixtures of monovalent and divalent cations. Proc. Natl. Acad. Sci. USA 2019, 116, 21022–21030.

(40) Kjellander, R. Modified Debye-Hückel approximation with effective charges: An application of dressed ion theory for electrolyte solutions. J. Phys. Chem. 1995, 99, 10392–10407.

(41) Frusawa, H.; Hayakawa, R. Generalizing the Debye-Hückel equation in terms of density functional integral. Phys. Rev. E 2000, 61, R6079–R6082.

(42) Denesyuk, N. A.; Thirumalai, D. Coarse-grained model for predicting RNA folding thermodynamics. J. Phys. Chem. B 2013, 117, 4901–4911.

(43) Schaefer, M.; Karplus, M. A comprehensive analytical treatment of continuum electrostatics. J. Phys. Chem. 1996, 100, 1578–1599.

(44) Bashford, D.; Case, D. A. Generalized born models of macromolecular solvation effects. Annu. Rev. Phys. Chem. 2000, 51, 129–152.

(45) Murthy, C. S.; Bacquet, R. J.; Rossky, P. J. Ionic distributions near polyelectrolytes. A comparison of theoretical approaches. J. Phys. Chem. 1985, 89, 701–710.

(46) Netz, R. R.; Orland, H. Beyond Poisson-Boltzmann: Fluctuation effects and correlation functions. Eur. Phys. J. E 2000, 1, 203–214.

(47) Attard, P. Electrolytes and the electric double layer. Adv. Chem. Phys. 1996, 92, 1–159.

(48) Shi, Y. Z.; Wang, F. H.; Wu, Y. Y.; Tan, Z. J. A coarse-grained model with implicit salt for RNAs: Predicting 3D structure, stability and salt effect. J. Chem. Phys. 2014, 141, 105102 1–13.

(49) Marcovitz, A.; Levy, Y. Frustration in protein-DNA binding influences conformational switching and target search kinetics. Proc. Natl. Acad. Sci. U. S. A. 2011, 108, 17957– 17962.

(50) Givaty, O.; Levy, Y. Protein sliding along DNA: Dynamics and structural characterization. J. Mol. Biol. 2009, 385, 1087–1097.

(51) Wong, G. C.; Pollack, L. Electrostatics of strongly charged biological polymers: Ion-mediated interactions and self-organization in nucleic acids and proteins. Annu. Rev. Phys. Chem. 2010, 61, 171–189.

(52) Savelyev, A.; Papoian, G. A. Molecular renormalization group coarse-graining of electrolyte solutions: Application to aqueous NaCl and KCl. J. Phys. Chem. B 2009, 113, 7785–7793.

(53) Savelyev, A.; Papoian, G. A. Chemically accurate coarse graining of double-stranded DNA. Proc. Natl. Acad. Sci. USA 2010, 107, 20340–20345.

(54) Denesyuk, N. A.; Hori, N.; Thirumalai, D. Molecular simulations of ion effects on the thermodynamics of RNA folding. J. Phys. Chem. B 2018, 122, 11860–11867.

(55) Åqvist, J. Ion-water interaction potentials derived from free energy perturbation simulations. J. Phys. Chem. 1990, 94, 8021–8024.

(56) Joung, I. S.; Cheatham, III, T.E. Determination of alkali and halide monovalent ion parameters for use in explicitly solvated biomolecular simulations. J. Phys. Chem. B 2008, 112, 9020–9041.

(57) Chen, A. A.; Pappu, R. V. Parameters of monovalent ions in the AMBER-99 forcefield:Assessment of inaccuracies and proposed improvements. J. Phys. Chem. B 2007, 111, 11884–11887.

(58) Kirmizialtin, S.; Pabit, S. A.; Meisburger, S. P.; Pollack, L.; Elber, R. RNA and its ionic cloud: Solution scattering experiments and atomically detailed simulations. Biophys. J. 2012, 102, 819–828.

(59) Chen, A. A.; García, A. E. High-resolution reversible folding of hyperstable RNA tetraloops using molecular dynamics simulations. Proc. Natl. Acad. Sci. USA 2013, 110, 16820–16825.

(60) Miner, J. C.; García, A. E. Concentration-dependent and configuration-dependent interactions of monovalent ions with an RNA tetraloop. J. Chem. Phys. 2018, 148, 222837 1–11.

(61) Srivastava, A.; Timsina, R.; Heo, S.; Dewage, S. W.; Kirmizialtin, S.; Qiu, X. Structure-guided DNA–DNA attraction mediated by divalent cations. Nucleic Acid Res. 2020, 48, 7018–7026.

(62) Arenz, S.; Bock, L. V.; Graf, M.; Innis, C. A.; Beckmann, R.; Grubmüller, H.; Vaiana, A. C.; Wilson, D. N. A combined cryo-EM and molecular dynamics approach reveals the mechanism of ErmBL-mediated translation arrest. Nat. Commun. 2016, 7, 1–14.

(63) Whitford, P. C.; Noel, J. K.; Gosavi, S.; Schug, A.; Sanbonmatsu, K. Y.; Onuchic, J. N. An all-atom structure-based potential for proteins: Bridging minimal models with allatom empirical forcefields. Prot. Struct. Func. Bioinfo. 2009, 75, 430–441.

(64) Noel, J. K.; Levi, M.; Raghunathan, M.; Lammert, H.; Hayes, R. L.; Onuchic, J. N.; Whitford, P. C. SMOG 2: A versatile software package for generating structure-based models. PLoS Comput. Biol. 2016, 12, e1004794 1–14.

(65) Lindorff-Larsen, K.; Piana, S.; Palmo, K.; Maragakis, P.; Klepeis, J. L.; Dror, R. O.; Shaw, D. E. Improved side-chain torsion potentials for the Amber ff99SB protein force field. Prot. Struct. Func. Bioinfo. 2010, 78, 1950–1958.

(66) Whitford, P. C.; Jiang, W.; Serwer, P. Simulations of phage T7 capsid expansion reveal the role of molecular sterics on dynamics. Viruses 2020, 12, 1273 1–14.

(67) Noel, J. K.; Whitford, P. C.; Onuchic, J. N. The shadow map: A general contact definition for capturing the dynamics of biomolecular folding and function. J. Phys. Chem. B 2012, 116, 8692–8702.

(68) Zhang, C.; Raugei, S.; Eisenberg, B.; Carloni, P. Molecular dynamics in physiological solutions: Force fields, alkali metal ions, and ionic strength. J. Chem. Theory Comput. 2010, 6, 2167–2175.

(69) Timasheff, S. N. Water as ligand: Preferential binding and exclusion of denaturants in protein unfolding. Biochem. 1992, 31, 9857–9864.

(70) Eisenberg, H. Biological macromolecules and polyelectrolytes in solution; Clarendon Press, 1976.

(71) Rice, S. A.; Nagasawa, M. Polyelectrolyte Solutions; Academic Press: New York, 1961.

(72) Grilley, D.; Misra, V.; Caliskan, G.; Draper, D.E. Importance of partially unfolded conformations for Mg2+-induced folding of RNA tertiary structure: Structural models and free energies of Mg2+ interactions. Biochem. 2007, 46, 10266–10278.

(73) Leipply, D.; Draper, D. E. Effects of Mg^2+^ on the free energy landscape for folding a purine riboswitch RNA. Biochem. 2011, 50, 2790–2799.

(74) Serganov, A.; Yuan, Y.-R.; Pikovskaya, O.; Polonskaia, A.; Malinina, L.; Phan, A. T.; Hobartner, C.; Micura, R.; Breaker, R. R.; Patel, D. J. Structural basis for discriminative regulation of gene expression by adenine- and guanine-sensing mRNAs. Chem. Biol. 2004, 11, 1729–1741.

(75) Noeske, J.; Schwalbe, H.; Wöhnert, J. Metal-ion binding and metal-ion induced folding of the adenine-sensing riboswitch aptamer domain. Nuc. Acids Res. 2007, 35, 5262– 5273.

(76) Conn, G. L.; Gittis, A. G.; Lattman, E. E.; Misra, V. K.; Draper, D. E. A compact RNA tertiary structure contains a buried backbone–K^+^ complex. J. Mol. Biol. 2002, 318, 963–973.

(77) Hayes, R. L.; Noel, J. K.; Mohanty, U.; Whitford, P. C.; Hennelly, S. P.; Onuchic, J. N.; Sanbonmatsu, K. Y. Magnesium fluctuations modulate RNA dynamics in the SAM-I riboswitch. J. Am. Chem. Soc. 2012, 134, 12043–12053.

(78) Hayes, R. L.; Noel, J. K.; Whitford, P. C.; Mohanty, U.; Sanbonmatsu, K. Y.; Onuchic, J. N. Reduced model captures Mg^2+^-RNA interaction free energy of riboswitches. Biophys. J. 2014, 106, 1508–1519.

(79) Misra, V. K.; Draper, D. E. A thermodynamic framework for Mg^2+^ binding to RNA. Proc. Natl. Acad. Sci. USA 2001, 98, 12456–12461.

(80) Xi, K.; Wang, F. H.; Xiong, G.; Zhang, Z. L.; Tan, Z. J. Competitive binding of Mg^2+^ and Na^+^ ions to nucleic acids: from helices to tertiary structures. Biophys. J. 2018, 114, 1776–1790.

(81) Lammert, H.; Wang, A.; Mohanty, U.; Onuchic, J. N. RNA as a complex polymer with coupled dynamics of ions and water in the outer solvation sphere. J. Phys. Chem. B 2018, 122, 11218–11227.

(82) Rodnina, M. V.; Wintermeyer, W. Fidelity of aminoacyl-tRNA selection on the ribosome: kinetic and structural mechanisms. Annu. Rev. Biochem. 2001, 70, 415–435.

(83) Whitford, P. C.; Geggier, P.; Altman, R. B.; Blanchard, S. C.; Onuchic, J. N.; Sanbonmatsu, K. Y. Accommodation of aminoacyl-tRNA into the ribosome involves reversible excursions along multiple pathways. RNA 2010, 16, 1196–1204.

(84) Noel, J. K.; Chahine, J.; Leite, V. B. P.; Whitford, P. C. Capturing Transition Paths and Transition States for Conformational Rearrangements in the Ribosome. Biophys. J. 2014, 107, 2872–2881.

(85) Yang, H.; Noel, J. K.; Whitford, P. C. Anisotropic Fluctuations in the Ribosome Determine tRNA Kinetics. J. Phys. Chem. B 2017, 121, 2777–2786.

(86) Yang, H.; Perrier, J.; Whitford, P. C. Disorder guides domain rearrangement in elongation factor Tu. Prot. Struct. Func. Bioinfo. 2018, 86, 1037–1046.

(87) Noel, J. K.; Whitford, P. C. How EF-Tu can contribute to efficient proofreading of aa-tRNA by the ribosome. Nat. Commun. 2016, 7, 13314.

(88) Whitford, P.; Onuchic, J.; Sanbonmatsu, K. Connecting Energy Landscapes with Experimental Rates for Aminoacyl-tRNA Accommodation in the Ribosome. J. Am. Chem. Soc. 2010, 132, 13170–13171.

(89) Yang, H.; Bandarkar, P.; Horne, R.; Leite, V. B.; Chahine, J.; Whitford, P. C. Diffusion of tRNA inside the ribosome is position-dependent. J. Chem. Phys. 2019, 151, 085102.

(90) Sanbonmatsu, K. Y.; Joseph, S.; Tung, C.-S. Simulating movement of tRNA into the ribosome during decoding. Proc. Natl. Acad. Sci. USA 2005, 102, 15854–15859.

(91) Rozov, A.; Khusainov, I.; El Omari, K.; Duman, R.; Mykhaylyk, V.; Yusupov, M.; Westhof, E.; Wagner, A.; Yusupova, G. Importance of potassium ions for ribosome structure and function revealed by long-wavelength X-ray diffraction. Nat. Commun. 2019, 10, 1–12.

(92) Johansson, M.; Lovmar, M.; Ehrenberg, M. Rate and accuracy of bacterial protein synthesis revisited. Curr. Opin. Microbio. 2008, 11, 141–147.

(93) Conn, G. L.; Draper, D. E.; Lattman, E. E.; Gittis, A. G. Crystal structure of a conserved ribosomal protein-RNA complex. Science 1999, 284, 1171–1174.

(94) Agrawal, R. K.; Linde, J.; Sengupta, J.; Nierhaus, K. H.; Frank, J. Localization of L11 protein on the ribosome and elucidation of its involvement in EF-G-dependent translocation. J. Mol. Biol. 2001, 311, 777–787.

(95) Gao, H.; Sengupta, J.; Valle, M.; Korostelev, A.; Eswar, N.; Stagg, S. M.; Van Roey, P.; Agrawal, R. K.; Harvey, S. C.; Sali, A.; Chapman, M. S.; Frank, J. Study of the structural dynamics of the E. coli 70S ribosome using real-space refinement. Cell 2003, 113, 789–801.

(96) Valle, M.; Zavialov, A.; Li, W.; Stagg, S. M.; Sengupta, J.; Nielsen, R. C.; Nissen, P.; Harvey, S. C.; Ehrenberg, M.; Frank, J. Incorporation of aminoacyl-tRNA into the ribosome as seen by cryo-electron microscopy. Nat. Struct. Biol. 2003, 10, 899–906.

(97) Seo, H.-S.; Abedin, S.; Kamp, D.; Wilson, D. N.; Nierhaus, K. H.; Cooperman, B. S. EF-G-dependent GTPase on the ribosome. Conformational change and fusidic acid inhibition. Biochem. 2006, 45, 2504–2514.

(98) Rodnina, M. V.; Savelsbergh, A.; Matassova, N. B.; Katunin, V. I.; Semenkov, Y. P.; Wintermeyer, W. Thiostrepton inhibits the turnover but not the GTPase of elongation factor G on the ribosome. Proc. Natl. Acad. Sci. USA 1999, 96, 9586–9590.

(99) Hayes, R. L.; Noel, J. K.; Mandic, A.; Whitford, P. C.; Sanbonmatsu, K. Y.; Mohanty, U.; Onuchic, J. N. Generalized Manning condensation model captures the RNA ion atmosphere. Phys. Rev. Lett. 2015, 114, 258105 1–6.

(100) Hyeon, C.; Thirumalai, D. Capturing the essence of folding and functions of biomolecules using coarse-grained models. Nat. Commun. 2011, 2, 487.

(101) Oliveira, A. B.; Contessoto, V. G.; Hassan, A.; Byju, S.; Wang, A.; Wang, Y.; Dodero-Rojas, E.; Mohanty, U.; Noel, J. K.; Onuchic, J. N.; Whitford, P. C. SMOG 2 and OpenSMOG: Expanding the limiting of structure-based models. Protein Science 2022, 31, 158–172.

(102) Nguyen, K.; Yang, H.; Whitford, P. C. How the ribosomal A-site finger can lead to tRNA species-dependent dynamics. J. Phys. Chem. B 2017, 121, 2767–2775.

(103) Levi, M.; Walak, K.; Wang, A.; Mohanty, U.; Whitford, P. C. A steric gate controls hybrid-state formation of tRNA on the ribosome. Nat. Commun. 2020, 11, 1–12.

(104) Trobro, S.; Åqvist, J. Role of ribosomal protein L27 in peptidyl transfer. Biochemistry 2008, 47, 4898–4906.

(105) Vaiana, A. C.; Sanbonmatsu, K. Y. Stochastic gating and drug-ribosome interactions. J Mol Biol 2009, 386, 648–661.

(106) Trabuco, L. G.; Schreiner, E.; Eargle, J.; Cornish, P.; Ha, T.; Luthey-Schulten, Z.; Schulten, K. The role of L1 stalk-tRNA interaction in the ribosome elongation cycle. J. Mol. Biol. 2010, 402, 741–760.

(107) Bock, L. V.; Blau, C.; Schröder, G. F.; Davydov, I. I.; Fischer, N.; Stark, H.; Rodnina, M. V.; Vaiana, A. C.; Grubmüller, H. Energy barriers and driving forces in tRNA translocation through the ribosome. Nat. Struct. Mol. Biol. 2013, 20, 1390–1396.

(108) Johansson, M.; Ieong, K. W.; Trobro, S.; Strazewski, P.; Åqvist, J.; Pavlov, M. Y.; Ehrenberg, M. pH-sensitivity of the ribosomal peptidyl transfer reaction dependent on the identity of the A-site aminoacyl-tRNA. Proc. Natl. Acad. Sci. USA 2011, 108, 79–84.

(109) Bock, L. V.; Blau, C.; Vaiana, A. C.; Grubmüller, H. Dynamic contact network between ribosomal subunits enables rapid large-scale rotation during spontaneous translocation. Nucleic Acid Res. 2015, 43, 6747–6760.

(110) Sethi, A.; Eargle, J.; Black, A. A.; Luthey-Schulten, Z. Dynamical networks in tRNA:protein complexes. Proc. Natl. Acad. Sci. USA 2009, 106, 6620–6625.

(111) Eargle, J.; Black, A. A.; Sethi, A.; Trabuco, L. G.; Luthey-Schulten, Z. Dynamics of Recognition between tRNA and elongation factor Tu. J Mol Biol 2008, 377, 1382– 1405.

(112) Trobro, S.; Åqvist, J. Mechanism of peptide bond synthesis on the ribosome. Proc. Natl. Acad. Sci. USA 2005, 102, 12396–12400.

(113) Gindulyte, A.; Bashan, A.; Agmon, I.; Massa, L.; Yonath, A.; Karle, J. The transition state for formation of the peptide bond in the ribosome. Proc. Natl. Acad. Sci. USA 2007, 103, 13327–13332.

