## Supplemental Information for "Diffuse ions coordinate dynamics in a ribonucleoprotein assembly"

### Contents


**1 Supporting Methods** **3**

**2 Supporting Results** **12**

**3 Supporting Figures** **17**

### 1 Supporting Methods

#### Parameter refinement of effective potentials for diffuse ion interactions

The parameters in the effective potential $V_{\mathrm{E}}$ (Eq.3 in main text) were initially set using explicit-solvent simulations as a reference. From explicit-solvent simulations, we calculated the radial distribution functions (RDF, Fig. S4) for different types of ion-RNA, ion-ion and ion-protein interactions and then fit our potential to the PMF in order to provide initial estimates. Then, an iterative protocol was applied to refine the values of parameters in the effective potential. The details are described below.

##### Initial parameters

Initial values for the parameters in the effective potential $V_{\mathrm{E}}$ $(A, B^{(k)}, C^{(k)}, R^{(k)})$ defined in Eq. 3 were determined by fitting the functional form of the effective potential to the potentials of mean force (PMF) (Fig. S4b) for pairwise interactions (e.g. K-K, K-Cl, K-Mg, Mg-Cl, etc.), calculated from all-atom explicit-solvent MD simulations (Fig. S4a). Each interaction was decomposed into a coulomb term, an effective excluded volume repulsive term and Gaussian terms that account for ionic solvation shells. First, we subtracted out the electrostatic potential (Debye-Hückel) from the PMF and then fit the remaining potential energy terms ($V_{\mathrm{ion-excl}}$ and $V_{\mathrm{sol}}$) to the residuals. The coefficient A in $V_{\mathrm{sol}}$ was initially set by fitting the function $\frac{A}{r^{12}}$ to the short distance region of the PMF associated with the corresponding interaction (Fig. S4c, dashed blue line). In the functional form of $V_{\mathrm{sol}}$, the width ($C^{(k)}$), location ($R^{(k)}$) and amplitude ($B^{(k)}$) of each Gaussian were obtained by fitting to a corresponding wells and peaks of the PMF. The widths and positions of the Gaussian functions indicate the ranges and locations of solvation shells around each type of ion, and these values remain fixed during all additional parameter refinement steps. With $R^{(k)}$ and $C^{(k)}$ determined for the Gaussian-based interactions, $V_{\mathrm{E}}$ can be expressed as a linear combination of $\sum \frac{1}{r_{ij}^{12}}$ and $\sum e^{-C^{(k)}[r_{ij}-R^{(k)}]^2}$ with coefficients $A$ and $B^{(k)}$. These linear coefficients are then refined using the protocol described below.

Through this procedure, we obtained the initial estimates of parameters for ion-ion, ion-RNA and ion-protein interactions from explicit solvent simulations of 100 mM KCl and 10 mM $MgCl_2$ in water, helix 44 (h44, Fig. 1) from rRNA in 100 mM KCl and 10 mM $MgCl_2$ solution and ribosomal protein S6 (Fig. S3) in 100 mM KCl and 10 mM $MgCl_2$ solution respectively.

**Iterative parameter refinement protocol**

Starting from the initial guess of parameters $A$ and $B^{(k)}$ in Eq. 3, we performed an iterative parameter refinement strategy, where explicit-solvent simulations are used as a benchmark. While the complete protocol is described elsewhere [15], we provide an overview here, for completeness. This protocol allows for refinement of all parameters that are linear in the Hamiltonian. In this protocol, one considers a Hamiltonian of the form

$$H_{\mathrm{R}} = \sum_{\alpha=1}^{N} K_\alpha S_\alpha, \tag{S1}$$

where a set of $N$ physical observables are denoted by $S_\alpha$ with associated weights $K_\alpha$. In our model, $K_\alpha$ corresponds to the parameters $A$ (in $V_{\mathrm{ion-excl}}$) and $B^{(k)}$ (in $V_{\mathrm{sol}}$), while $S_\alpha$ corresponds to $\sum_{i<j} \frac{1}{r_{ij}^{12}}$ and $\sum_{i<j} e^{-C^{(k)}[r_{ij}-R^{(k)}]^2}$. As described in the previous section, $C^{(k)}$ and $R^{(k)}$ remain fixed during refinement.

The form of the Hamiltonian is exploited to construct an iterative scheme to determine the coefficient $K_\alpha$ associated with each observable $S_\alpha$ [9, 15, 16]. Let the expected value of the observable $S_\alpha$ be denoted by $\langle S_\alpha \rangle$. Thus, $\langle S_\alpha \rangle_{\mathrm{SBM}}$ and $\langle S_\alpha \rangle_{\mathrm{ex-sol}}$ denote the average values of the observables, as calculated from the SBM simulations and all-atom explicit-solvent simulation, respectively. To determine values of $S_\alpha$ that accurately describe observables in a reference system, we expand the expectation value of each observable $\langle S_\alpha \rangle_{\mathrm{SBM}}$ in a Taylor series in $K_\alpha$ around some trial weights $\{K_\alpha^{(0)}\}$. The difference between the expectation values of the observables, $\Delta\langle S_\alpha\rangle = \langle S_\alpha\rangle_{\rm SBM} - \langle S_\alpha\rangle_{\rm ex-sol}$, to first-order is given by [9, 15, 16]

$$\Delta\langle S_\alpha\rangle = \sum_\gamma \frac{\partial\langle S_\alpha\rangle}{\partial K_\gamma}\Delta K_\gamma \tag{S2a}$$

$$= \frac{1}{k_B T}\sum_\gamma [\langle S_\alpha S_\gamma\rangle - \langle S_\alpha\rangle\langle S_\gamma\rangle]\Delta K_\gamma, \tag{S2b}$$

where $\Delta K_\gamma^{(n)} = K_\gamma^{(n+1)} - K_\gamma^{(n)}$. Eq. (S2b) follows from Eq. (S2a) due to the linearity of the reference Hamiltonian.

To obtain the corrections to the weights, $\Delta K_\alpha^{(n)}$, we iteratively performed SMOG-ion simulations and update the weights in each iteration by solving the set of linear equations, i.e., Eq. (S2b). The *n*th iteration, for example, defines the parameter set $K_\alpha^{(n+1)} = K_\alpha^{(n)} + \Delta K_\alpha^{(n)}$. The iterative simulation and updates of the weights are continued until $K_\alpha$ converges for $\alpha = 1, ..., N$. As $K_\alpha$ converges, the observable $S_\alpha$ obtained from the refined model converges to that from the reference system and the RDF from SMOG-ion model simulation converges to that from the explicit solvent simulation (Fig. S13a). The values of coefficients $A$ and $B^{(k)}$ are obtained from the converged values of $K_\alpha$.

As employed in previous studies conducted by Savelyev and Papoian [15], the quality of the parameters can be measured by calculating the difference in free-energy ($|\delta F|$) between the explicit-solvent and structure-based model, defined as:

$$\delta F = \sum_\alpha |K_\alpha \Delta S_\alpha| \tag{S3}$$

In the iterative simulations, $\delta F$ decreased significantly in the first few iterations and then levels off near zero (Fig. S13). In the parameter refinement process, we stopped updating parameters when $\delta F$ reached a minimal value.

#### Defining non-contact interactions in the SMOG-ion model

Any atom pairs that are not in contact in the predefined structure are defined as "non-contact interactions". In the SMOG-ion model, the non-contact interaction of a pair of atoms (i, j) in RNA or protein that are separated by $r_{ij}$ have a potential energy of the form $\frac{C_{18}}{r_{ij}^{18}} - \frac{C_{12}}{r_{ij}^{12}}$. This pair-wise potential accounts for the soft repulsive interactions between atoms due to the excluded volume. In order to mimic the excluded volume description provided by all-atom explicit-solvent models, the coefficients $C_{18}$ and $C_{12}$ were calculated by fitting the 12-18 potential to the Lennard-Jones potential ($V_{\mathrm{Amber-LJ}}$) in the Amber99sb-ildn forcefield [8], which is given by $4\epsilon[(\frac{\sigma}{r_{ij}})^{12} - (\frac{\sigma}{r_{ij}})^{6}]$, where $\sigma$ and $\epsilon$ are provided in the force field parameter set. Two reference points, $(V_{\mathrm{Amber-LJ,1}}, r_{ij,1})$ and $(V_{\mathrm{Amber-LJ,2}}, r_{ij,2})$, are obtained for each pair of atoms from the Amber99sb-ildn forcefield [8], where $V_{\mathrm{Amber-LJ,1}} = 1/2k_BT$ and $V_{\mathrm{Amber-LJ,2}} = k_BT$. The value of coefficients $C_{18}$ and $C_{12}$ are calculated by fitting the functional form of the non-contact interaction to the reference points. An example of the curve fitting is shown in Figure S1.

#### Simulation details

##### *Explicit-solvent simulations*

Explicit-solvent MD simulations were used as a benchmark for initial parameterization of the SMOG-ion model. The explicit-solvent simulations were conducted using Gromacs v5.1.4 with the Amber99sb-ildn force field [8]. The simulated systems were solvated with SPC/E water molecules [2]. Modified $Mg^{2+}$ parameters described by Åqvist [1] were used in the simulation. The parameters for monovalent ions ($K^{+}$ and $Cl^{-}$) reported by Joung and Cheatham [6] were included. Periodic boundary conditions were used in the simulations. Particle-mesh Ewald (PME) was used for the evaluation of long-range electrostatics [4] with an Ewald radius of 10 Å. The Van der Waals cutoff was taken to be 10 Å. Neighbor lists were updated every 10 time steps. Equations of motion were integrated using a leap-frog integrator with a 2-fs time step.

In order to parameterize the ion-ion, ion-RNA and ion-protein interactions in the SMOG-ion model, the following explicit-solvent simulations were performed as benchmarks:

1. *Ions in aqueous solution.* 60 $K^+$, 6 $Mg^{2+}$, 72 $Cl^-$ ions and 32635 water molecules were added to a 10 nm cubic box to create a target concentration of 10 mM for $[MgCl_2]$ and 100 mM for [KCl]. The water molecules and ions were energetically minimized with steepest descent followed by conjugate gradient methods. The system was initially equilibrated at 300 K and ambient pressure of 1 bar using the Berendsen thermostat and barostat [3] for 1 ns. After the equilibration step, $1\mu$s of production runs was carried out with Nose-Hoover thermostat [5, 11] and Parrinello-Rahman barostat [12, 14]. This simulation provides benchmark reference for ion-ion interactions.

2. *Helix 44 in ionic solution.* A RNA fragment, helix 44 (h44), from 16S ribosomal RNA (PDB: 4V6F) was placed in the center of a $10 \times 12.2 \times 12.2$ nm$^3$ rectangular box with the principal axis of H44 aligned with the x-dimension of the box. 123 $K^+$, 14 $Mg^{2+}$, 108 $Cl^-$ and 47917 water molecules were added to the box to neutralize the negative charges from H44 and yield a bulk concentrations of approximately 10 mM for $[MgCl_2]$ and 100 mM for [KCl]. To eliminate the influence from any conformational changes of h44 on ion-RNA interactions, every atom on H44 was position restrained to its initial coordinate with force constant 1000 kJ/mol/nm$^2$. The entire system was energetically minimized with steepest descent and then conjugate gradient method. Then we performed the equilibration of the system at 300 K and 1 bar with Berendsen thermostat and barostat [3], which was followed by 1 $\mu$s production run with Nose-Hoover thermostat [5, 11] and Parrinello-Rahman barostat [12, 14]. The first 20 ns trajectory was used for equilibrating ions with the h44 and was excluded from RDF analysis. This simulation provides the benchmark reference for ion-RNA interactions.

3. *Protein S6 in ionic solution.* Protein S6 was extracted from a structure of a bacterial ribosome

(PDB: 4V6F) and was placed in the center of a rectangular box of size $10 \times 16 \times 16\text{nm}^3$ with its first principal axis aligned to the x-dimension of the box. 15 $Mg^{2+}$, 154 $K^+$ and 185 $Cl^-$ ions were added to the box together with 88364 water molecules to create a bulk concentration of approximately 10 mM $[MgCl_2]$ and 100 mM of [KCl]. Atoms in the protein molecule were position restrained to their initial coordinates with force constant 1000 kJ/mol/nm$^2$ during the simulations. The energy of the system was minimized with steepest descent and conjugate gradient methods. The entire system was equilibrated at 300 K and 1 bar with Berendsen thermostat and barostats [3], which was followed by the 1 $\mu$s production run with Nose-Hoover thermostat [5, 11] and Parrinello-Rahman barostat [12, 14]. The first 20 ns trajectory was excluded from the RDF analysis. This simulation provides the benchmark reference for ion-protein interactions.

***SMOG-ion model simulations***

MD simulations with the SMOG-ion model were performed for different purposes: (1) refining the parameters A and B in the effective potential $V_{\mathrm{E}}$ (Eq. 3) in comparison with explicit-solvent simulations, (2) investigating the preferential interaction coefficients ($\Gamma_{2+}$) of two RNA molecules with refined parameters to compare with experimental measurements, (3) studying the conformational dynamics of the ribosome structure under ionic effects. Simulation details are as followed.

1. *SMOG-ion model simulation during parameterization.* For the purpose of parameter refinement, SMOG-ion simulations were performed with exactly the same number of ions in the same size of boxes as in the corresponding benchmark explicit-solvent simulations. The simulations were performed at a temperature of 60 Gromacs units (0.5 reduced units) which corresponds to 300 K in explicit-solvent simulations. In each iteration of parameter refinement, 10 replicas of 10 million timesteps were performed with the SMOG-ion model, and trajectories from all replicas were used to determine the value of each observable.

2. *SMOG-ion model simulations for small RNA molecules.* After the parameter refinement process, SMOG-ion simulations were applied to the small 58-mer rRNA (PDB: 1HC8) and the adenine riboswitch (PDB: 1Y26) to investigate the ion preferential interaction coefficient ($\Gamma_{2+}$) for each system. In both cases, the small RNA molecule was placed in a cubic box of length 70 nm. Ions were placed randomly but not close to each other or the RNA molecule initially. For the 58mer rRNA, the number of $Mg^{2+}$ ions was varied from 23 to 216 to achieve the target bulk concentration of $Mg^{2+}$ ions (0.10-1.00 mM). 30983 $K^{+}$ ions were placed in the box to set the concentration of $K^{+}$ to 150mM. For the adenine riboswitch, 23 to 207 $Mg^{2+}$ ions were used with 10328 $K^{+}$ ions to create a $Mg^{2+}$ concentrations that range of 0.088 to 1.00 mM and a $K^{+}$ concentration of 50 mM. In both cases, $Cl^{-}$ ions were added to neutralize the charges in the system. The chelated ions were harmonically bound to the neighboring oxygen atoms in the chelation pocket. At each concentration, the system was equilibrated for $10^8$ timesteps with 0.002 reduced unit for each timestep. Then the equilibrium run was extended for another $2 \times 10^7$ and we took a snapshot of the atom coordinates after each $10^6$ timesteps. These 20 snapshots were used to initialize 20 replicas with random initial velocities. Each replica was then simulated for an additional $6 \times 10^7$ time steps, from which the first $10^7$ time steps were discarded during analysis.

3. *SMOG-ion model simulations for ribosome.* The simulation of 70S ribosome is performed using a crystal structure (PDB: 6QNR) of the ribosome complex. In SMOG-ion model, the potential energy minimun of the ribosome structure is defined by the A/A conformation. A 4 Å cutoff was used to defined the native contacts [10]. To avoid unphysical expansion of the ribosome molecule, all contact distances in the ribosome were reduced by a factor of 0.96. In the process of aa-tRNA accommodation, since the intermolecular arm and 3'-CCA contacts only form after the elbow accommodation [17], they are not included in the simulations with SMOG-ion model and in the reference simulations with SMOG model (no diffuse ions

and no electrostatics). This modification was made such that the calculations in this work could specifically focus on the role of diffuse ions in the elbow accommodation step, not in the subsequent rearrangements. In addition, the stabilizing contacts between A-site and P-site tRNAs are reduced by a factor of 0.15, such that the free energy od the A/T and EA ensemble is comparable when the electrostatic-free model (SMOG model) is used. To obtain the initial A/T configuration of the aa-tRNA, a crystal strucutre (PDB: 4V5D) of ribosome bound to EF-Tu and aa-tRNA is used as a reference for aligning P-site tRNA and providing target positions of aa-tRNA in A/T configuration. All simulations of the ribosome were performed starting from the A/T configuration. In each simulation, the ribosome structure was placed in the center of a cubic box with length 70 nm. Different numbers (Tab. S5) of $Mg^{2+}$, $K^{+}$ and $Cl^{-}$ ions were added to the box to create different bulk concentrations of $Mg^{2+}$ from 0 to 2.67 mM while maintaing the bulk monovalent ion concentration as 100 mM. At each concentration, we ran $1.5 \times 10^7$ timesteps of equilibration in which the ribosome was position restrained at its initial coordinates, which allowed the diffuse ions to equilibrate around the ribosome. Then production runs were performed starting from the equilibrated system in various independent replicas. In simulations at $Mg^{2+}$ concentrations 0.1 and 0.27 mM, $1.7 \times 10^9$ timesteps were accumulated from 10 replicas. In simulations at $Mg^{2+}$ concentrations 0 mM and 2.67 mM, $6.0 \times 10^8$ timesteps were accumulated from 4 replicas.

***Software and model access***

Upon publication, all models described in this study will be freely available through the smog-server.org webpage. This includes a modified version of Gromacs 5.1.4 that supports the ion effective potentials, as well as the SMOG2 template files for the SMOG-ion model and the SMOG model with 12-18 excluded volume interactions. These force fields are also compatible with the OpenS-

MOG libraries [13] for use with OpenMM. The force field template files will be accessible through the smog-server Force Field repository page, with accession codes, `AA_ions_Wang21.v1` (SMOG-ion), `AA_12-18_Wang21.v1` (electrostatics-free SMOG with 12-18 non-bonded interactions). Additionally, the same force fields are available with cutoff contact maps, with accession codes `AA_ions_cutoff_Wang21.v1` and `AA_12-18_cutoff_Wang21.v1`.

### 2 Supporting Results

#### Refined effective potential parameters

As described in the Method section, the SMOG-ion model parameters were initially refined in comparison with explicit-solvent simulations. Then a few parameters of ion-RNA interactions were further adjusted by comparing with the experimental results and using the preferential interaction coefficient ($\Gamma_{2+}$) as a metric. The finalized parameter set s3 is shown in the tables below.

Table S1: Refined parameters for ion-ion interactions are obtained from the simulation with 10mM $MgCl_2$ and 100mM KCl. $\epsilon = 2k_BT$ in the units of parameter A and B.

| Interaction | A $[\epsilon \cdot \mathrm{nm}^{12}]$ | B $[\epsilon]$ | | | | |
|---|---|---|---|---|---|---|
| | A | $B_1$ | $B_2$ | $B_3$ | $B_4$ | $B_5$ |
| $K^+$-$K^+$ | $2.510 \times 10^{-6}$ | -0.7371 | 0.1887 | -0.0029 | 0.111 | 0.0632 |
| $K^+$-$Cl^-$ | $4.484 \times 10^{-7}$ | -0.4908 | 0.7674 | -0.2481 | 0.0578 | -0.1116 |
| $K^+$-$Mg^{2+}$ | $1.639 \times 10^{-4}$ | -0.032 | 0.2484 | 0.0448 | 0.1482 | 0.1106 |
| $Cl^-$-$Cl^-$ | $4.689 \times 10^{-5}$ | -0.3862 | 0.348 | -0.0205 | 0.1262 | 0.0509 |
| $Mg^{2+}$-$Cl^-$ | $1.213 \times 10^{-5}$ | -0.4291 | 0.4266 | -0.2472 | -0.0106 | -0.1742 |
| $Mg^{2+}$-$Mg^{2+}$ | $9.215 \times 10^{-3}$ | - | - | - | - | - |

| Interaction | C $[\mathrm{nm}^{-2}]$ | | | | | R [nm] | | | | |
|---|---|---|---|---|---|---|---|---|---|---|
| | $C_1$ | $C_2$ | $C_3$ | $C_4$ | $C_5$ | $R_1$ | $R_2$ | $R_3$ | $R_4$ | $R_5$ |
| $K^+$-$K^+$ | 284.3 | 1039 | 1458.5 | 427.8 | 235.5 | 0.424 | 0.5662 | 0.6673 | 0.7712 | 0.8986 |
| $K^+$-$Cl^-$ | 1125 | 304.3 | 571.8 | 1309.9 | 392.1 | 0.3247 | 0.3924 | 0.5386 | 0.6358 | 0.7441 |
| $K^+$-$Mg^{2+}$ | 1769.9 | 204.7 | 335.6 | 216.2 | 1815.1 | 0.5933 | 0.6973 | 0.8095 | 0.9056 | 1.0144 |
| $Cl^-$-$Cl^-$ | 770.8 | 299.9 | 1844 | 514.2 | 650.7 | 0.5245 | 0.638 | 0.768 | 0.8631 | 0.984 |
| $Mg^{2+}$-$Cl^-$ | 628.3 | 798.3 | 359.2 | 2041.6 | 404.1 | 0.4715 | 0.5465 | 0.6775 | 0.7775 | 0.8794 |
| $Mg^{2+}$-$Mg^{2+}$ | - | - | - | - | - | - | - | - | - | - |

Table S2: Refined parameters for ion-RNA interaction. Atoms from RNA are grouped by their element and the value of their partial charges. Only the interactions between metal ions ($Mg^{2+}$ and $K^{+}$) with the negatively charged RNA atoms are parameterized. The negatively charged RNA atoms are grouped by their elements and the value of their partial charges. For atoms whose partial charges are less than -0.49e, the element symbols are superscripted with "<-0.5", otherwise, the element symbol is superscripted with ">-0.5". In the table, "0" means the corresponding parameters are defined to be zero, while "-" indicates the corresponding parameters are not defined in the SMOG-ion model.

| Interaction | A [$\epsilon \cdot \mathrm{nm}^{12}$] | B [$\epsilon$] | | | | |
|---|---|---|---|---|---|---|
| | A | $B_1$ | $B_2$ | $B_3$ | $B_4$ | $B_5$ |
| $K^{+}$-$O_{<-0.5}$ | $2.923 \times 10^{-8}$ | 0 | 0.2962 | 0.0339 | -0.0217 | -0.0973 |
| $K^{+}$-$N_{<-0.5}$ | $3.703 \times 10^{-8}$ | 0 | 0.085 | -0.2878 | - | - |
| $K^{+}$-$O_{>-0.5}$ | $3.688 \times 10^{-8}$ | 0 | 0.9727 | - | - | - |
| $K^{+}$-$C_{>-0.5}$ | $1.208 \times 10^{-6}$ | - | - | - | - | - |
| $K^{+}$-$N_{>-0.5}$ | $2.239 \times 10^{-7}$ | - | - | - | - | - |
| $Mg^{2+}$-$O_{<-0.5}$ | $2.522 \times 10^{-6}$ | -0.89 | 0.0157 | -0.1161 | 0.0093 | -0.0599 |
| $Mg^{2+}$-$N_{<-0.5}$ | $2.250 \times 10^{-6}$ | -1.02 | - | - | - | - |
| $Mg^{2+}$-$O_{>-0.5}$ | $1.465 \times 10^{-5}$ | - | - | - | - | - |
| $Mg^{2+}$-$C_{>-0.5}$ | $1.600 \times 10^{-5}$ | - | - | - | - | - |
| $Mg^{2+}$-$N_{>-0.5}$ | $1.162 \times 10^{-5}$ | - | - | - | - | - |

| Interaction | C [$\mathrm{nm}^{-2}$] | | | | | R [nm] | | | | |
|---|---|---|---|---|---|---|---|---|---|---|
| | $C_1$ | $C_2$ | $C_3$ | $C_4$ | $C_5$ | $R_1$ | $R_2$ | $R_3$ | $R_4$ | $R_5$ |
| $K^{+}$-$O_{<-0.5}$ | 1095 | 501 | 438 | 1833 | 123 | 0.278 | 0.348 | 0.511 | 0.569 | 0.698 |
| $K^{+}$-$N_{<-0.5}$ | 1067 | 325 | 180 | - | - | 0.286 | 0.37 | 0.617 | - | - |
| $K^{+}$-$O_{>-0.5}$ | 954 | 255 | - | - | - | 0.285 | 0.38 | - | - | - |
| $K^{+}$-$C_{>-0.5}$ | - | - | - | - | - | - | - | - | - | - |
| $K^{+}$-$N_{>-0.5}$ | - | - | - | - | - | - | - | - | - | - |
| $Mg^{2+}$-$O_{<-0.5}$ | 651 | 392 | 331 | 305 | 92.7 | 0.417 | 0.478 | 0.61 | 0.725 | 1.01 |
| $Mg^{2+}$-$N_{<-0.5}$ | 646 | - | - | - | - | 0.441 | - | - | - | - |
| $Mg^{2+}$-$O_{>-0.5}$ | - | - | - | - | - | - | - | - | - | - |
| $Mg^{2+}$-$C_{>-0.5}$ | - | - | - | - | - | - | - | - | - | - |
| $Mg^{2+}$-$N_{>-0.5}$ | - | - | - | - | - | - | - | - | - | - |

Table S3: Refined parameter for ion-protein interaction. Atoms in the protein are also grouped by element and their assigned partial charges. Repulsive interactions between ion and protein atoms are parameterized with both excluded volume and Gaussian terms. For the attractive interactions, only excluded volume coefficients were refined. If an atom has a more negative charge than -0.49e, a subscription "<-0.5" will be added to the element symbol. If an atom has a charge between -0.49e and 0, a subscription ">-0.5" will be added to the element symbol. If an atom has positive partial charge in between - and 0.49e, its element symbol has a subscription "<0.5". If an atom has partial charge greater than 0.49e, its element symbol has a subscription ">0.5". There are a few special cases in this table: "$C_{<0}$" consists of both "$C_{<-0.5}$" and "$C_{>-0.5}$" atoms; "$N_{<0}$" consists of both "$N_{<-0.5}$" and "$N_{>-0.5}$" atoms.

| Interaction | A [$\epsilon \cdot nm^{12}$] | B [$\epsilon$] | | | | |
|---|---|---|---|---|---|---|
| | A | $B_1$ | $B_2$ | $B_3$ | $B_4$ | $B_5$ |
| $K^+$-$O_{<-0.5}$ | $2.898 \times 10^{-8}$ | -1.1804 | 0.2875 | -0.0972 | - | - |
| $K^+$-$O_{>-0.5}$ | $7.092 \times 10^{-8}$ | -1.2356 | 0.371 | -0.0959 | - | - |
| $K^+$-$N_{<0}$ | $1.515 \times 10^{-6}$ | - | - | - | - | - |
| $K^+$-$N_{>0.5}$ | $3.340 \times 10^{-6}$ | - | - | - | - | - |
| $K^+$-$N_{<0.5}$ | $3.756 \times 10^{-7}$ | - | - | - | - | - |
| $K^+$-$C_{<0}$ | $1.635 \times 10^{-6}$ | - | - | - | - | - |
| $K^+$-$S_{>-0.5}$ | $4.297 \times 10^{-6}$ | - | - | - | - | - |
| $Mg^{2+}$-$O_{<-0.5}$ | $7.160 \times 10^{-7}$ | -1.3849 | 0.1479 | -0.0034 | - | - |
| $Mg^{2+}$-$O_{>-0.5}$ | $2.345 \times 10^{-6}$ | - | - | - | - | - |
| $Mg^{2+}$-$N_{<0}$ | $8.740 \times 10^{-5}$ | - | - | - | - | - |
| $Mg^{2+}$-$N_{>0.5}$ | $7.638 \times 10^{-4}$ | - | - | - | - | - |
| $Mg^{2+}$-$N_{<0.5}$ | $1.764 \times 10^{-5}$ | - | - | - | - | - |
| $Mg^{2+}$-$C_{<0}$ | $1.852 \times 10^{-5}$ | - | - | - | - | - |
| $Mg^{2+}$-$S_{>-0.5}$ | $9.425 \times 10^{-5}$ | - | - | - | - | - |
| $Cl^-$-$N_{>0.5}$ | $5.504 \times 10^{-7}$ | -0.944 | 0.1997 | -0.3535 | -0.0562 | -0.1387 |
| $Cl^-$-$N_{<0.5}$ | $6.004 \times 10^{-7}$ | -0.9644 | 0.0828 | -0.1301 | - | - |
| $Cl^-$-$C_{>0.5}$ | $4.707 \times 10^{-6}$ | - | - | - | - | - |
| $Cl^-$-$C_{<0.5}$ | $2.323 \times 10^{-6}$ | - | - | - | - | - |
| $Cl^-$-$C_{<0}$ | $2.420 \times 10^{-6}$ | - | - | - | - | - |
| $Cl^-$-$O_{<-0.5}$ | $8.554 \times 10^{-6}$ | - | - | - | - | - |
| $Cl^-$-$S_{>-0.5}$ | $1.193 \times 10^{-5}$ | - | - | - | - | - |

Table S4: Refined parameter for ion-protein interaction (continued).

| Interaction | C [$nm^{-2}$] | | | | | R [nm] | | | | |
|---|---|---|---|---|---|---|---|---|---|---|
| | $C_1$ | $C_2$ | $C_3$ | $C_4$ | $C_5$ | $R_1$ | $R_2$ | $R_3$ | $R_4$ | $R_5$ |
| $K^+$-$O_{<-0.5}$ | 1127 | 376 | 481 | - | - | 0.27 | 0.351 | 0.498 | - | - |
| $K^+$-$O_{>-0.5}$ | 535 | 186 | 444 | - | - | 0.272 | 0.381 | 0.542 | - | - |
| $K^+$-$N_{<0}$ | - | - | - | - | - | - | - | - | - | - |
| $K^+$-$N_{>0.5}$ | - | - | - | - | - | - | - | - | - | - |
| $K^+$-$N_{<0.5}$ | - | - | - | - | - | - | - | - | - | - |
| $K^+$-$C_{<0}$ | - | - | - | - | - | - | - | - | - | - |
| $K^+$-$S_{>-0.5}$ | - | - | - | - | - | - | - | - | - | - |
| $Mg^{2+}$-$O_{>-0.5}$ | 524 | 1158 | 418 | - | - | 0.406 | 0.477 | 0.656 | - | - |
| $Mg^{2+}$-$O_{>-0.5}$ | - | - | - | - | - | - | - | - | - | - |
| $Mg^{2+}$-$N_{<0}$ | - | - | - | - | - | - | - | - | - | - |
| $Mg^{2+}$-$N_{>0.5}$ | - | - | - | - | - | - | - | - | - | - |
| $Mg^{2+}$-$N_{<0.5}$ | - | - | - | - | - | - | - | - | - | - |
| $Mg^{2+}$-$C_{<0}$ | - | - | - | - | - | - | - | - | - | - |
| $Mg^{2+}$-$S_{>-0.5}$ | - | - | - | - | - | - | - | - | - | - |
| $Cl^-$-$N_{>0.5}$ | 1298 | 253 | 284 | 1719 | 307 | 0.337 | 0.395 | 0.546 | 0.652 | 0.758 |
| $Cl^-$-$N_{<0.5}$ | 1098 | 298 | 1130 | - | - | 0.338 | 0.395 | 0.558 | - | - |
| $Cl^-$-$C_{>0.5}$ | - | - | - | - | - | - | - | - | - | - |
| $Cl^-$-$C_{<0.5}$ | - | - | - | - | - | - | - | - | - | - |
| $Cl^-$-$C_{<0}$ | - | - | - | - | - | - | - | - | - | - |
| $Cl^-$-$O_{<-0.5}$ | - | - | - | - | - | - | - | - | - | - |
| $Cl^-$-$S_{>-0.5}$ | - | - | - | - | - | - | - | - | - | - |

Table S5: Ionic conditions of ribosome simulations

| model name | num. diffuse $Mg^{2+}$ | num. diffuse $K^{+}$ | num. diffuse $Cl^{-}$ | bulk [$Mg^{2+}$] (mM) |
|---|---|---|---|---|
| SMOG (no charges) | 0 | 0 | 0 | 0 |
| SMOG-ion | 0 | 23000 | 20226 | 0 |
| SMOG-ion | 300 | 22000 | 19826 | 0.11 |
| SMOG-ion | 500 | 22000 | 20226 | 0.27 |
| SMOG-ion | 1500 | 20000 | 20226 | 2.67 |

### 3 Supporting Figures

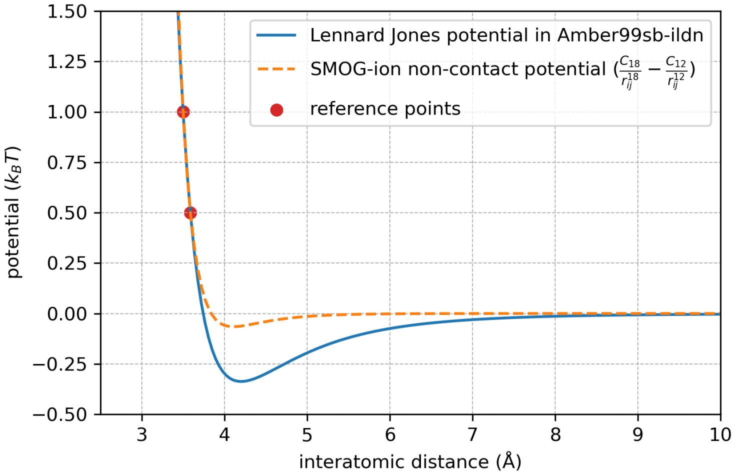


Figure S1: **Paramaterizing excluded volume interactions based on the AMBER force field.** An example for how the coefficients $C_{18}$ and $C_{12}$ were defined (orange dashed line) in the SMOG-ion model. The Lennard-Jones potential (blue solid curve) defined in Amber99sb-ildn forcefield [8] provides the reference values (red dots).

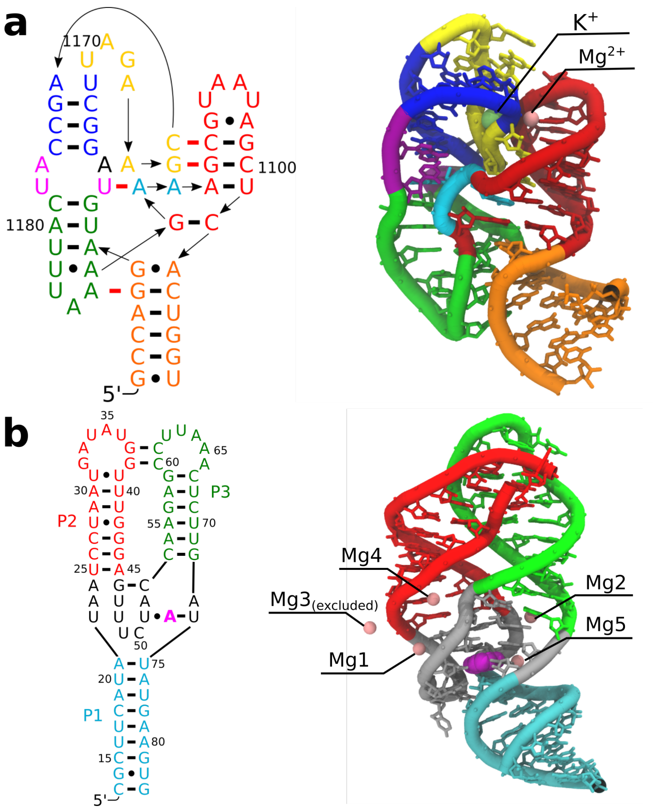


Figure S2: **Structures of 58-mer rRNA fragment and adenine riboswitch.** (a) 58-mer rRNA fragment (PDB:1HC8). Color codes of residues on right panel are consistent with the left panel. In the left panel, base pairs are indicated by black horizontal lines, tertiary base-base hydrogen bonds in the folded RNA are shown in red bars. Arrows indicate the 5' to 3' direction of the backbone. Two chelated ions ($K^+$ and $Mg^{2+}$) are shown in beads. (b) Adenine riboswitch (PDB:1Y26). Color codes of residues in the tertiary structure on right panel are consistent with the left panel. 5 chelated $Mg^{2+}$ ions are shown in pink.

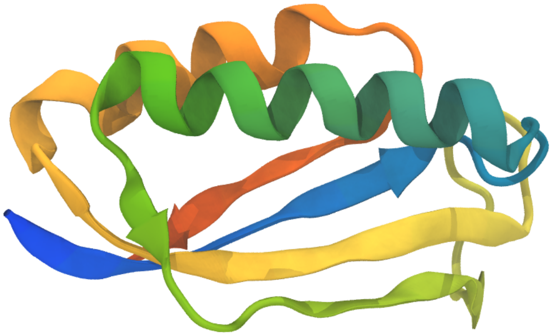

Figure S3: **Structure of protein S6**. S6 is a small protein with 101 amino acid residues. It consists two $\beta-\alpha-\beta$ motifs with four-stranded anti-parallel $\beta$-sheet on one side and two $\alpha$-helices packed on the other side. Similar folding patterns are observed in other ribosomal proteins. [7]. Since our goal is to investigate the effects of diffuse ions on ribonucleoprotein assemblies, we decided to use this globular ribosomal protein as a reference when parameterizing the interactions between ions and protein atoms.

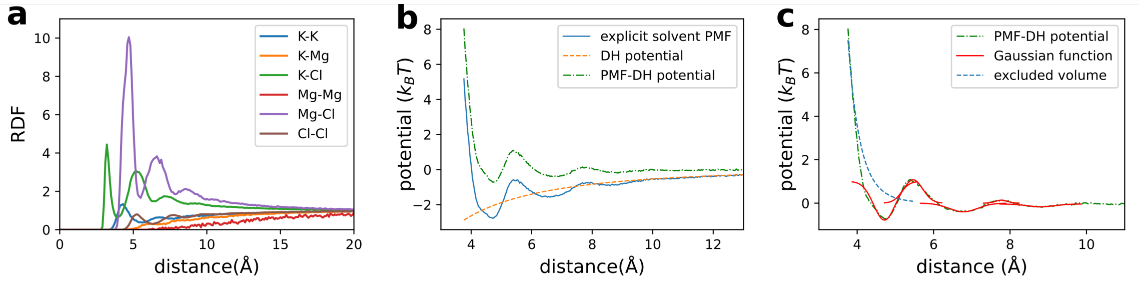


Figure S4: **Explicit-solvent simulations provide reference for ion association interactions.** (a) The RDFs of ion-ion interactions from explicit solvent simulations of 10 mM $MgCl_2$ and 100 mM KCl in water. (b) The PMF of Mg-Cl interactions (blue), which is converted from the corresponding RDF values. The PMF based on the Debye-Hückel (DH) potential is shown as dashed line (orange). The PMF with DH contribution subtracted is shown as dashed dot line (green). (c) For initial fitting of the parameters in the Hamiltonian (see Eq.3) that describe Mg-Cl interactions, the DH contribution was subtracted from the PMF (dashed dot green line). The resulting PMF was approximated by the excluded volume repulsive term (dashed blue line) and the sum of five Gaussian functions (red line).

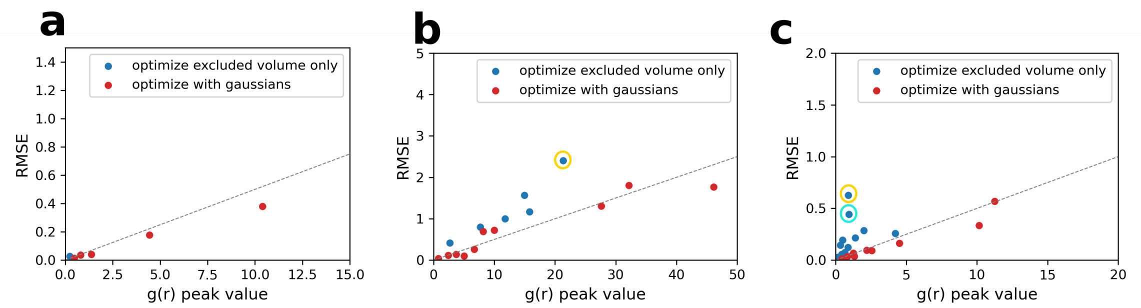


Figure S5: **Consistency of parameterized interactions in the SMOG-ion model and an explicit-solvent model.** Root mean squared error (RMSE) of g(r) between SMOG-ion model (s1 parameter set, see main text for definition) simulation and the corresponding explicit solvent simulation versus the peak g(r) values of the latter are plotted for each type of interactions. Dashed line in each panel corresponds to the boundary of 5% relative error. Generally, interactions optimized with Gaussians in their effective potentials (red dots) benefit from the parameterization of the solvation effects, which results in lower relative error comparing to those optimized for excluded volume only (blue dots). a) The RMSE associated with ion-ion interactions from the simulations of ions in aqueous solution. b) The RMSE associated with ion-RNA and ion-ion interactions from the simulations of H44 in ionic solution. The blue dot in the yellow circle, which associates with the interaction between $Mg^{2+}$ ion and weakly negatively charged C atoms ($C_{>-0.5}$) in RNA residues, yields the highest relative error (11.2%) in this set. All other interactions have relative error within 10%. c) The RMSE associated with ion-protein and ion-ion interactions from the simulation of protein S6 in ionic solution. The interaction between $Mg^{2+}$ ion and weakly positively charged N atoms ($N_{<0.5}$, yellow circle) yield the highest relative error, because $N_{<0.5}$ only appears in ARG amino acid residues in protein S6 and has nearly zero (0.0329 $e$) charge, which result super weak interactions with $Mg^{2+}$ ions and obvious fluctuations in g(r). The second highest relative error associates with the interaction between $Mg^{2+}$ ion and the highly positively charged N atom ($N_{>0.5}$), because the latter only appears in N-terminal amino acid residues and LYS residue, which only appears 4 times in protein S6 and results insufficient samplings for precise parameterization.

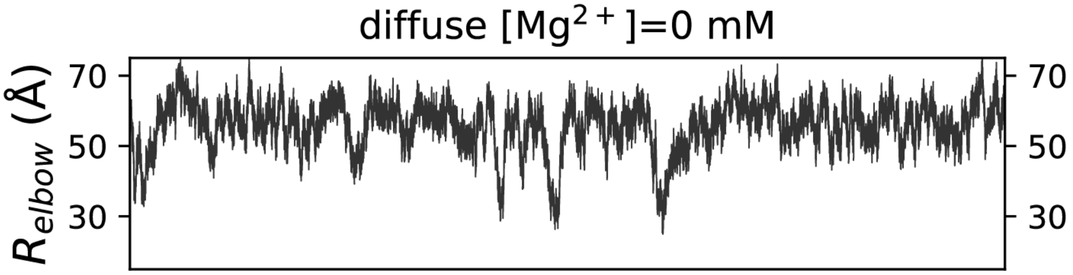


Figure S6: **Representative trajectory of SMOG-ion simulations of aa-tRNA accommodation in the presence of 100 mM $K^+$ and no diffuse $Mg^{2+}$ ions.**

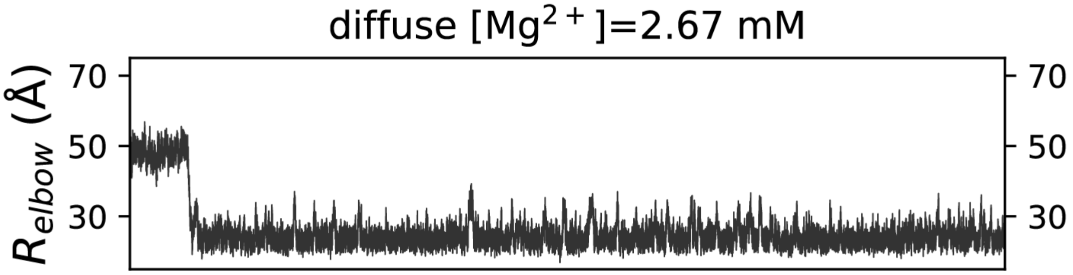


Figure S7: **Representative trajectory of SMOG-ion simulations of aa-tRNA accommodation in the presence of 100 mM $K^+$ and 2.67 mM $Mg^{2+}$ ions.**

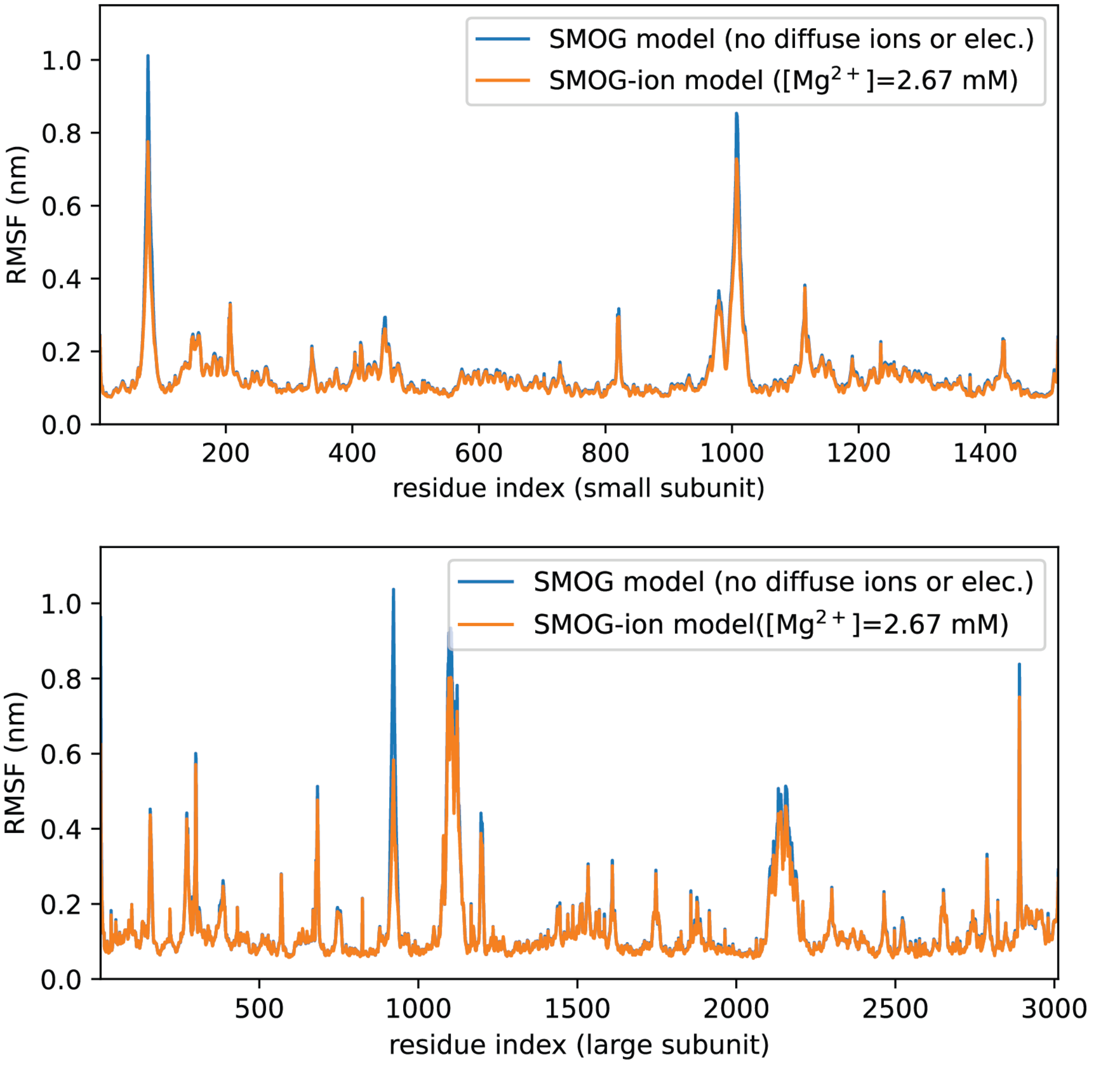


Figure S8: **Root mean squared fluctuations (RMSF) of rRNA from simulations.** RMSF is calculated for the rRNA in the ribosome and averaged for each residue.

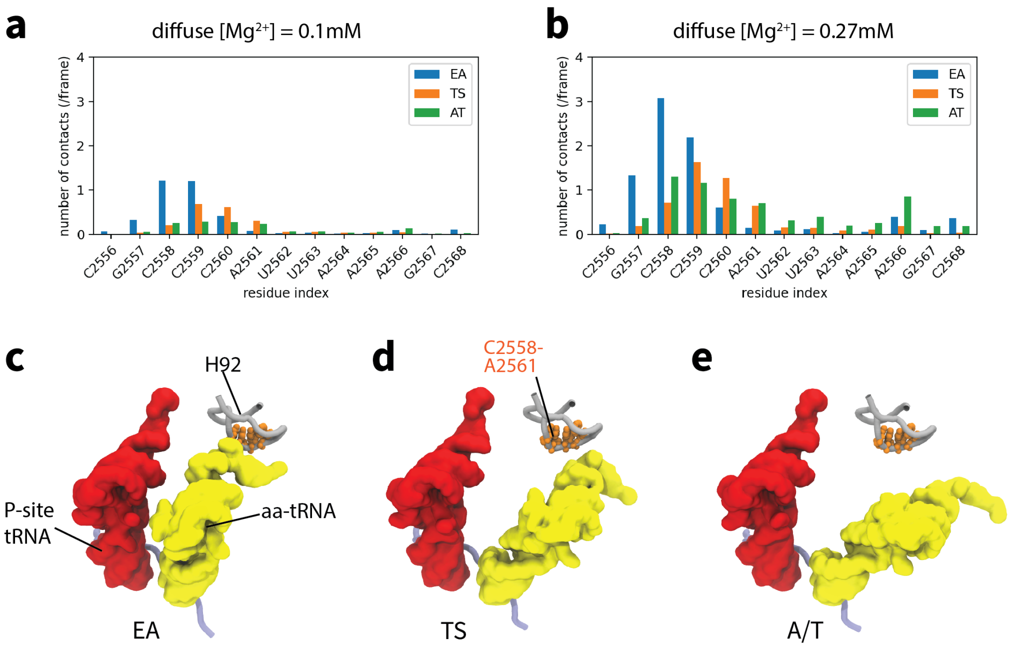


Figure S9: **$Mg^{2+}$-mediated interactions between Helix 92 (H92) and aa-tRNA stabilizes EA configuration.** a) Number of $Mg^{2+}$-mediated interactions between aa-tRNA and H92 when the bulk concentration of $Mg^{2+}$ is 0.1 mM. b) Number of $Mg^{2+}$-mediated interactions between aa-tRNA and H92 when the bulk concentration of $Mg^{2+}$ is 0.27 mM. c) Representative structure of H92 with aa-tRNA in EA state. d) Representative structure of H92 with aa-tRNA in TSE. e) Representative structure of H92 with aa-tRNA in A/T state.

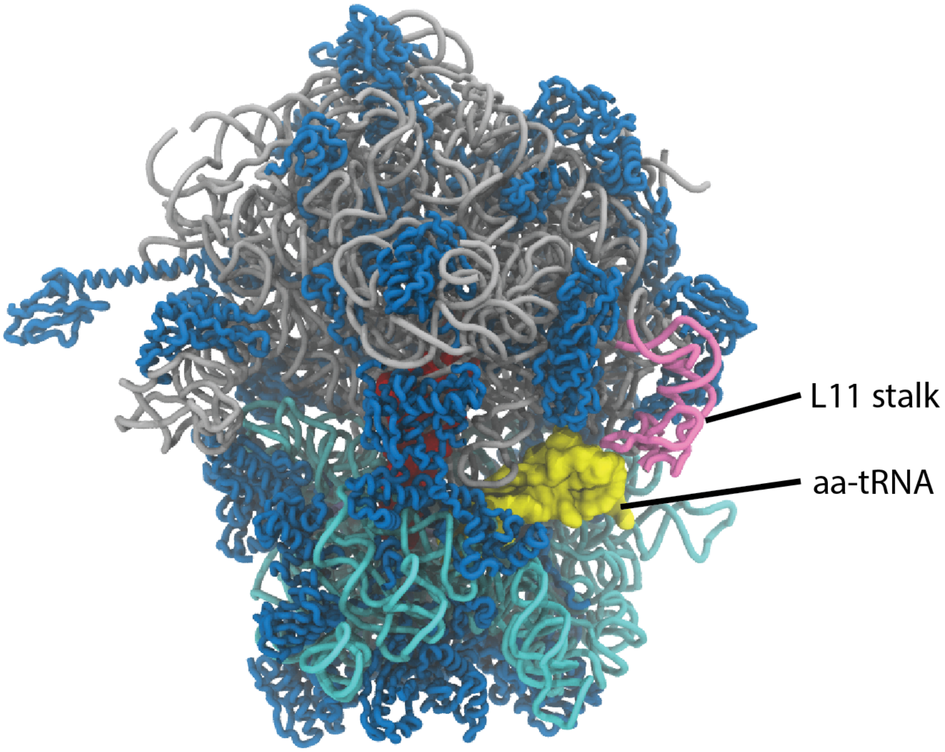


Figure S10: **L11 stalk (pink) is located at the periphery of the ribosome.**

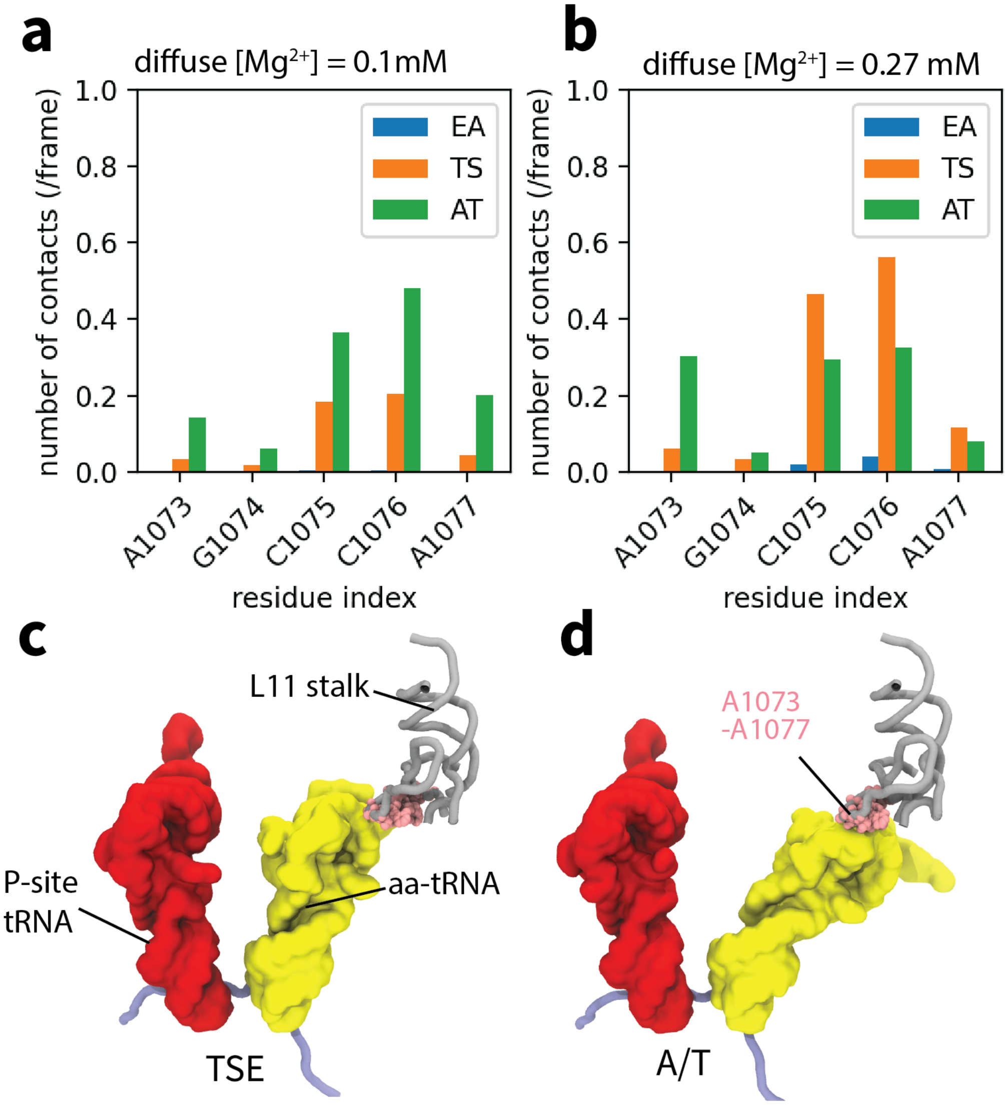


Figure S11: **$Mg^{2+}$-mediated interactions between L11 stalk and aa-tRNA stabilizes A/T configuration.** a) Number of $Mg^{2+}$-mediated interactions between aa-tRNA and L11 stalk when the bulk concentration of $Mg^{2+}$ is 0.1 mM. b) Number of $Mg^{2+}$-mediated interactions between aa-tRNA and L11 stalk when the bulk concentration of $Mg^{2+}$ is 0.27 mM. c) Representative structure of L11 stalk with aa-tRNA in TSE. d) Representative structure of L11 stalk with aa-tRNA in A/T state.

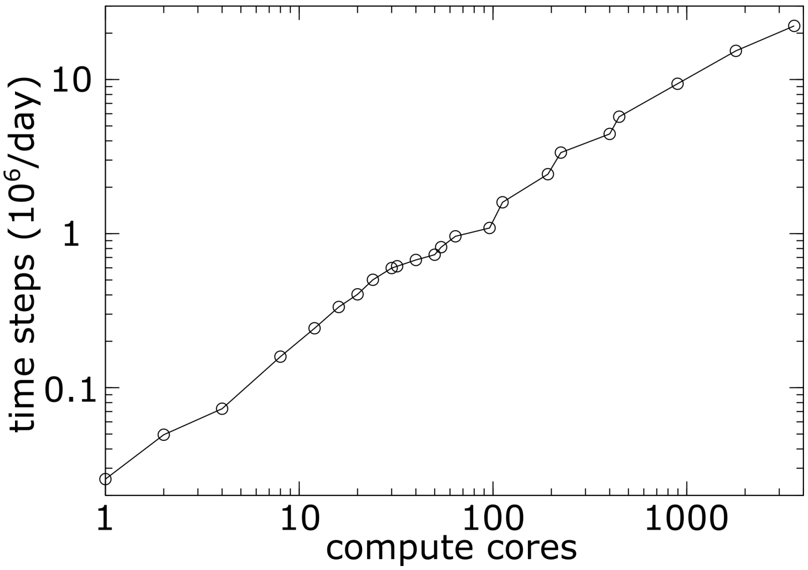


Figure S12: **Simulation performance as a function of the number of compute cores utilized.** Scaling tests were performed for the ribosome with diffuse ions (198,654 atoms). Performance gains were obtained for up to 3,584 compute cores. Tests used Gromacs 5.1.4, with modified non-bonded kernels to accommodate the SMOG-ion model. Each compute node was equipped with two Intel Xeon Platinum 8276 processors (2.20GHz) with Infiniband interconnect between nodes. Larger core counts were not tested due to the availability of resources.

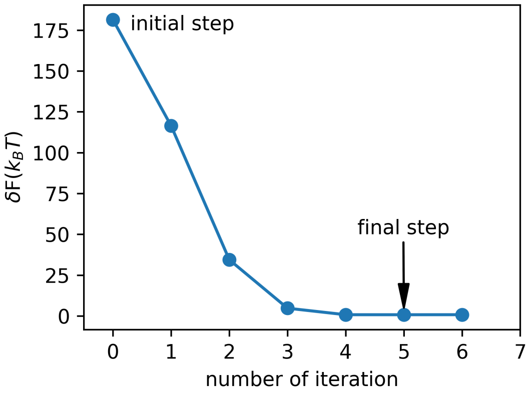


Figure S13: **Iterative parameter refinement leads to energetic consistency with explicit-solvent simulations.**The reduction of the total free energy difference ($\delta F$, Eq. S3) between explicit solvent model and structure-based model along the parameter refinement of ion-RNA interactions. The parameter refinement is terminated when $\delta F$ reaches a minimal value.

### References


[1] J. Åqvist. Ion-water interaction potentials derived from free energy perturbation simulations. *J. Phys. Chem.*, 94(21):8021–8024, oct 1990.

[2] H. J. Berendsen, J. R. Grigera, and T. P. Straatsma. The missing term in effective pair potentials. *J. Phys. Chem.*, 91(24):6269–6271, nov 1987.

[3] H. J. Berendsen, J. P. Postma, W. F. Van Gunsteren, A. Dinola, and J. R. Haak. Molecular dynamics with coupling to an external bath. *J. Phys. Chem.*, 81(8):3684–3690, 1984.

[4] T. Darden, D. York, and L. Pedersen. Particle mesh Ewald: An N·log(N) method for Ewald sums in large systems. *J. Chem. Phys.*, 98(12):10089–10092, jun 1993.

[5] W. G. Hoover. Canonical dynamics: Equilibrium phase-space distributions. *Phys. Rev. A*, 31(3):1695–1697, mar 1985.

[6] I. S. Joung and T. E. Cheatham, III. Determination of alkali and halide monovalent ion parameters for use in explicitly solvated biomolecular simulations. *J. Phys. Chem. B*, 112(30):9020–9041, jul 2008.

[7] M. Lindahl, L. A. Svensson, A. Liljas, S. E. Sedelnikova, I. A. Eliseikina, N. P. Fomenkova, N. Nevskaya, S. V. Nikonov, M. B. Garber, T. A. Muranova, A. I. Rykonova, and R. Amons. Crystal structure of the ribosomal protein S6 from Thermus thermophilus. *EMBO J.*, 13(6):1249–1254, 1994.

[8] K. Lindorff-Larsen, S. Piana, K. Palmo, P. Maragakis, J. L. Klepeis, R. O. Dror, and D. E. Shaw. Improved side-chain torsion potentials for the Amber ff99SB protein force field. *Prot. Struct. Func. Bioinfo.*, 78(8):1950–1958, 2010.

[9] A. P. Lyubartsev and A. Laaksonen. Calculation of effective interaction potentials from radial distribution functions: A reverse Monte Carlo approach. *Phys. Rev. E*, 52(4):3730–3737, oct 1995.

[10] J. K. Noel, P. C. Whitford, and J. N. Onuchic. The shadow map: A general contact definition for capturing the dynamics of biomolecular folding and function. *J. Phys. Chem. B*, 116:8692–8702, 2012.

[11] S. Nosé. A unified formulation of the constant temperature molecular dynamics methods. *J. Chem. Phys.*, 81(1):511–519, 1984.

[12] S. Nosé and M. L. Klein. Constant pressure molecular dynamics for molecular systems. *Mol. Phys.*, 50(5):1055–1076, dec 1983.

[13] A. B. Oliveira, V. G. Contessoto, A. Hassan, S. Byju, A. Wang, Y. Wang, E. Dodero-Rojas, U. Mohanty, J. K. Noel, J. N. Onuchic, and P. C. Whitford. SMOG 2 and OpenSMOG: Expanding the limiting of structure-based models. *Protein Science*, 31:158–172, 2022.

[14] M. Parrinello and A. Rahman. Polymorphic transitions in single crystals: A new molecular dynamics method. *J. Appl. Phys.*, 52(12):7182–7190, dec 1981.

[15] A. Savelyev and G. A. Papoian. Molecular renormalization group coarse-graining of electrolyte solutions: Application to aqueous NaCl and KCl. *J. Phys. Chem. B*, 113(22):7785–7793, jun 2009.

[16] R. H. Swendsen. Monte Carlo Renormalization Group. *Phys. Rev. Lett.*, 42(14):859–861, apr 1979.

[17] P. C. Whitford, P. Geggier, R. B. Altman, S. C. Blanchard, J. N. Onuchic, and K. Y. Sanbonmatsu. Accommodation of aminoacyl-tRNA into the ribosome involves reversible excursions along multiple pathways. *RNA*, 16(6):1196–1204, jun 2010.
